# Causal necessity of human hippocampus for structure-based inference in learning

**DOI:** 10.1101/2025.08.19.664920

**Authors:** Deng Pan, Simone D’Ambrogio, Naomi Kingston, Miruna Rascu, Pranav Sankhe, Shuyi Luo, Miriam C. Klein-Flügge, Ali Mahmoodi, Matthew F.S. Rushworth

## Abstract

When meeting new individuals or encountering known individuals in new circumstances, we intuitively map out their relationships – not merely by direct experience, but by quickly inferring new connections based on prior relational knowledge. Using a novel task, we demonstrated that participants indeed employ knowledge of relational structures to facilitate learning of new relationships in a changing environment. Computational modelling revealed that participants leveraged relational knowledge to support inference, thus facilitating learning. Whole brain neuroimaging identified a uniquely robust representation of relational structure in the hippocampus. Neural networks trained on similar tasks demonstrated the emergence of relational structure representations, resembling those found in hippocampus. Lesioning network units sustaining such representations disrupted structure-based inference and predicted hippocampus’s essential role. Transcranial ultrasound stimulation of human hippocampus, transiently modulating its activity without affecting overlying tissue, produced similar disruption effects, empirically confirming the causal necessity of hippocampal representations for structure-based inference in learning.

## Introduction

We have an increasingly detailed understanding of the neural mechanisms that underlie learning and decision making; we understand how people and other animals learn about the values of options, how options are compared, and the neural circuits that mediate these processes^1–4^. In the real world, however, our ability to make flexible decisions in changing environments surpasses what might be expected if we had to learn about options anew each time circumstances change. This is because, as we experience how one thing has changed in a new environment, we make inferences about other things in that new environment even before experiencing them directly^5,6^. This ability is particularly salient in abstract and social domains, where we often intuitively extend knowledge from one relationship to infer others. However, it remains unknown how this is accomplished in the brain.

One possibility is that the brain can build mental models of the environment^7–9^ to inform our predictions in the face of changes or even reversals in circumstances. The hippocampus (HPC) has been implicated in mental models that represent not only spatial relations^10^, but also conceptual and social relations^11,12^. There is, however, particularly in humans, little evidence to link such hippocampal representations to decision-making in changing environments. In the current study, we show that HPC extracts the underlying relational structure from a dynamic environment to facilitate learning of new relations. More importantly, we further demonstrate that these representations are causally necessary for structure-based inference during new learning. This causal link is established by employing a novel non-invasive technique, focused transcranial ultrasound stimulation (TUS), to temporarily modulate bilateral HPC activity. As a deep brain region, HPC has been inaccessible to conventional non-invasive methods in humans^13^. TUS, however, can precisely, safely, and reversibly modulate deep brain activity, as demonstrated by recent work with animal models^14–21^, allowing us to assess the causal effects of hippocampal disruption.

Through a series of experiments, we empirically assess the possibility that learning in changing environments is facilitated by knowledge of relational structure. We find that this is indeed the case even in the simple world of an artificial game (Fig. 1). In the game, participants learned the relational structure between four characters and each character’s preference, both of which underwent reversals. When the same underlying relational structure is present across situations, this is reflected in HPC representations, and participants learn rapidly and adapt flexibly in new situations. Computational modelling reveals that such knowledge of relational structure facilitates learning by enabling structure-based inference beyond direct experience. Functional magnetic resonance imaging (fMRI) demonstrates a consistent representation of relational structure in HPC across diverse contexts, a property not observed in other brain regions. In contrast, frontal cortical regions, such as the orbitofrontal cortex (OFC), which primarily predict character preferences in the game, exhibit only intermittent relational representations when making these predictions.

**Figure 1.**
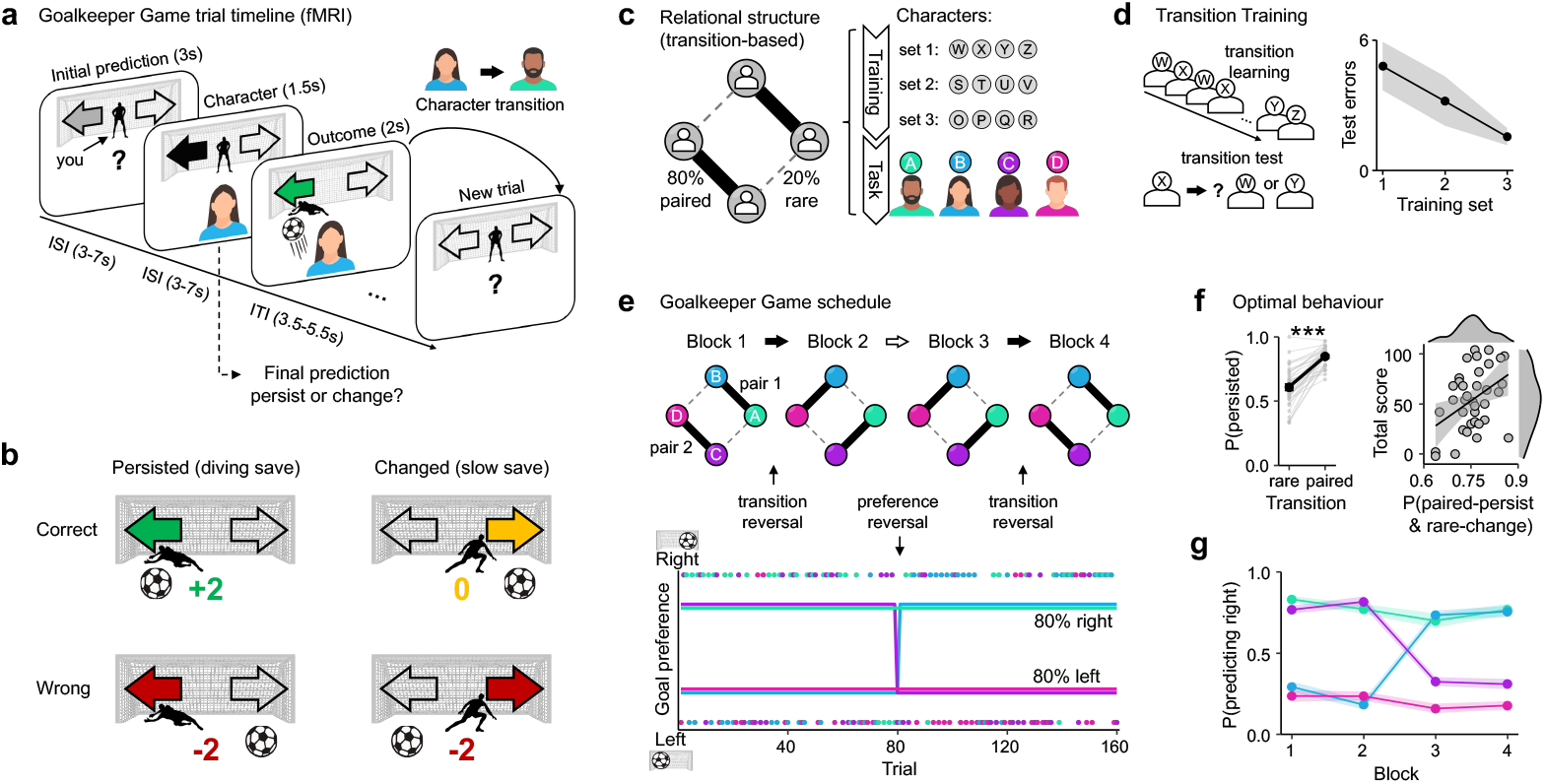
The experimental design and participants’ behaviours. **a**. Trial timeline of the Goalkeeper Game. In each trial, participants acted as goalkeepers, initially predicting the direction of the upcoming shot before knowing the character who would take the shot. After a jittered inter-stimulus interval (ISI), the character appeared on the screen. At that time, participants could decide to either persist with or change their initial prediction. The outcome, the actual goal direction, was displayed after another ISI as a football icon and a score earned on that trial (coloured arrow). Every time a new trial began, participants had to anticipate the next character based on the relational structure (transitions) between characters to make initial predictions. **b**. Payoff matrix. The score was based on a combination of prediction accuracy and whether participants persisted with or changed their initial predictions. Persist-correct (+2, green): participants persisted with the initial prediction and prediction was correct (“diving save”); Change-correct (0 points, yellow): participants changed the initial prediction during the character phase, and the new prediction was correct (“slow save”); In both ‘persist-wrong’ and ‘change-wrong’ scenarios (−2, red), participants were penalised −2 points for any wrong prediction, regardless of whether they persisted or changed. **c**. Transition-based relational structure. On a Markov chain, four characters are bidirectionally connected by transition probabilities, forming two pairs. Thick solid lines indicate paired transitions (80% probability), and thin dashed lines indicate rare transitions (20% probability). Both the training and the main task followed the same transition structure but were instantiated using different character sets. **d**. Transition training. Before the main task, participants completed three training blocks solely to learn the underlying structure of the character transitions. Each transition training block consisted of two parts: learning and testing. In the learning part, participants observed four characters repeatedly appearing on the screen following a probabilistic sequence. In the testing part, participants were tested on their understanding of the transitions by selecting the most likely character to follow a given one. They had to achieve 90% accuracy to pass each block; otherwise, the block had to be repeated. As participants progressed through the training blocks, the total number of test errors (across all attempts in each training block) decreased. Shaded areas represent standard errors. **e**. Illustration of transition and preference reversals in the Goalkeeper Game. Four new characters (A, B, C, D) were initially grouped into two pairs (80% transition probability). Transition reversals occurred after Blocks 1 and 3 (upper panel), swapping paired and rare transitions. A preference reversal occurred after Block 2 (bottom panel), in which two of the characters swapped their goal preferences. Colours indicate character identity, lines show preferred directions (80% of trials), and dots represent the actual goal directions of each trial. **f**. Participants persisted more during paired transitions than during rare transitions (left panel). Grey dots connected by lines represent each participant’s data. \*\*\**P*<0.001. The proportion of optimal behaviour (paired-persist & rare-change) is positively correlated with their total score (right panel). Fitted regression lines are plotted, and the shaded area reflects 95% confidence intervals (CIs). Distributions of the data are shown on the edges of the plot. **g**. Proportion of rightward final predictions across blocks. Each coloured line represents the proportion of rightward final predictions for a given character. Shaded areas denote standard errors of the mean (SEM).

Next, to guide our investigation of the causal necessity of HPC in inference-based learning, we used neural network modelling to predict the behavioural effects of disrupting relational structure representations. Specifically, we built a meta-reinforcement learning (meta-RL) recurrent neural network (RNN) to model the acquisition of relational knowledge through a simple learning process similar to that of human participants.

After training, the RNN developed units that represented relational structures, resembling the representations in human HPC. Critically, lesioning these HPC-like units selectively disrupted the network’s ability to make inferences from relational knowledge during learning, providing computational predictions for the causal role of HPC representations in inference. Consistent with this prediction, using a novel TUS protocol to stimulate the entire HPC, we selectively disrupted participants’ ability to learn the changing relations, particularly abolishing structure-based inference, confirming that HPC is causally necessary for flexible learning.

## Results

### Experimental task and behavioural results

In the fMRI experiment, before the main task, participants (*N*=36) first completed a *Transition Training* task outside the scanner to learn the transition-based relational structure (i.e., transition structure) among four characters. The training consisted of three sessions, each featuring a distinct set of characters. However, all training sets shared the same underlying structure: a Markov chain with four states (characters) forming two pairs (Fig. 1c). Transitions between paired characters (i.e., paired transitions) occurred with a high probability (80%), while transitions between unpaired characters were rare (20%) or absent (0%). All transitions were bidirectional, and self-transitions are not allowed. In each training block, they observed a four-character sequence governed by the two-pair transition structure and were then tested on it (Fig. 1d & S1a). Although the structure was never made explicit, participants implicitly learned it, as evidenced by a gradual decline in their cumulative test errors before passing each training set (one*-*way repeated ANOVA: *F*(2,70)=3.465, *P=*0.037; Fig. 1d). This echoes previous findings that animals and humans can learn the structure of a problem from examples to solve new ones more effectively^5,22^. The training phase prepared participants for the main task, in which relational structures were similar, but the specific relationships changed at times, requiring continuous adaptation.

After completing the *Transition Training*, participants performed the main task, the *Goalkeeper Game*, while undergoing fMRI. Playing the role of a goalkeeper, participants predicted upcoming shots by learning both the relational structure among four new characters (*A, B, C*, and *D*) and each character’s goal direction preference. At the beginning of each trial, except the first, participants made an initial prediction of the next shot (left or right) before knowing who was taking the shot, as in a free-kick situation (Fig. 1a & S1b). Accurate initial prediction required participants to anticipate the upcoming character based on trial-by-trial character transitions. Notably, they were never explicitly asked to predict the next character, ensuring this character anticipation remained an implicit process rather than another response-driven subgoal. This design allows us to dissociate relational structure learning from other goal-directed behaviours like predicting character preferences. Shortly after the initial prediction, the character was revealed. Once the character’s identity was revealed, participants could either persist with their initial prediction (choose the same direction) or change it (choose the other direction). Then, the outcome—the character’s actual goal direction and the points earned—was revealed. Participants were rewarded +2 points for persisting with a correct initial prediction, 0 points for a correct save after changing their initial prediction, and were penalised −2 points for any wrong predictions (Fig. 1b). This payoff scheme introduced a cost for changing and incentivised accurate initial predictions in the first place, which required learning both the character transitions and each character’s preference.

Crucially, character transitions in the *Goalkeeper Game* followed the same two-pair structure as in the *Transition Training*. Having implicitly learned the structure during training, participants could generalise it to the new characters in the main game. However, in the *Goalkeeper Game*, the transitions between characters were not fixed and changed at some points throughout the game (“transition reversal”). After each transition reversal, the previous paired transitions (80%) became rare transitions (20%) and *vice versa* (i.e., rare transitions became paired transitions), while the underlying two-pair structure remained unchanged (Fig. 1e). For example, if the initial structure contained *AB & CD* pairs, it would contain *AC & BD* pairs after a reversal. Additionally, each character had a preferred goal direction (left or right, 80% of the time), which also underwent an unsignalled reversal (“preference reversal”), where two characters swapped their preferences (Fig. 1e). Consequently, regardless of the reversal, the group always comprised two characters preferring left and two preferring right (Fig. 2a). Transition and preference reversals occurred at different points in the task, which enabled clear separation between transition learning and preference learning. The game consisted of four blocks, each containing around 40 trials. Unbeknownst to participants, transition reversals occurred twice, at the start of blocks 2 and 4, and a single preference reversal occurred at the start of block 3 (Fig. 1e & S2a). These reversal manipulations not only echo those commonly used to study flexible behaviour^23^, but also mirror real-life sports scenarios: two pairs of players form distinct attacking units early on, then later recombine into new formations to launch attacks from unexpected angles. Most importantly, these reversals created a dynamic environment, requiring participants to continuously learn character transitions and character preferences.

**Figure 2.**
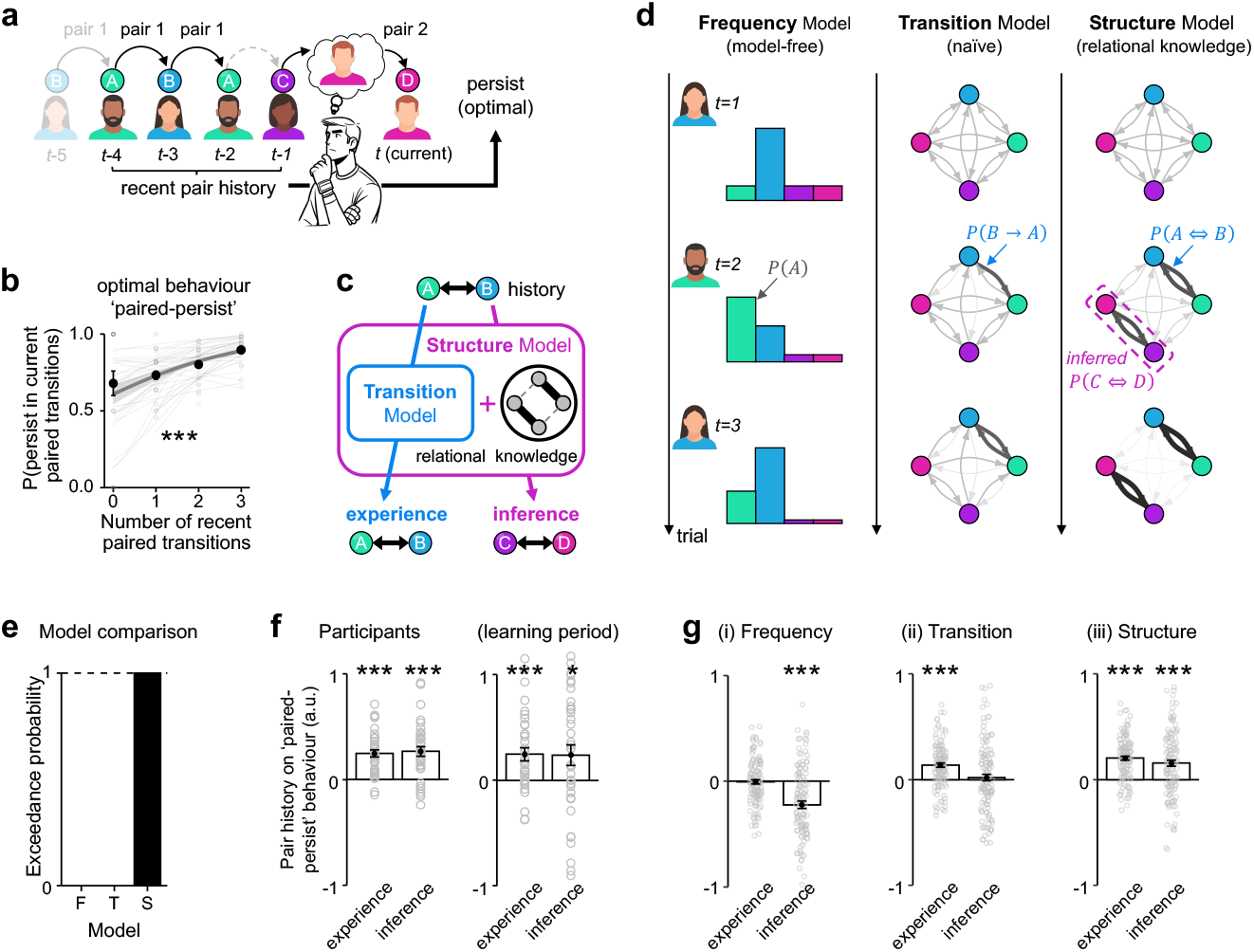
Computational modelling: relational knowledge supports inference-based learning. **a**. Illustration of how recent paired transition history can influence a participant’s decision to persist or change their initial prediction in a current paired transition (e.g., C-to-D, pair 2). In this example, the last three trials include two paired transitions (solid arrows) and one rare transition (dashed arrow). If the participant correctly anticipates D following C based on learned relational knowledge, the optimal behaviour is to persist with the initial prediction associated with D (“paired-persist”). **b**. The probability of participants persisting with their initial prediction increased with the number of recent paired transitions. Each grey dot represents one participant for a given paired transition history; thin light-grey lines show individual regression fits. Black dots with error bars show group mean ± SEM. The group-level regression fit is plotted as a thick, dark grey line with 95% CI (without random effect, for visualisation purposes only). **c**. The transition history can be further categorised into two types relative to the current transition. When recent history involves the same pair as the current pair, participants rely on direct experience for predictions. However, when recent history involves the other pair, they must rely on inference. While the Transition Model only learns from experience, the Structure Model extends it by incorporating relational knowledge, enabling inference beyond direct experience. **d**. Illustration of how each model learns sequences over time. In this example, two characters appear sequentially (B–A–B) from trial *t*=1 to *t*=3. The Frequency Model (Model F) tracks how often each character has appeared. The Transition Model (Model T) updates transition probabilities between characters based only on direct observations. The Structure Model (Model S) updates both observed and inferred transitions simultaneously by leveraging prior knowledge of the two-pair structure. The learning rates for all simulations are set to 0.5. **e**. The model comparison results show that the Structure Model (Model S) provides the best fit for the participants’ behavioural data, with an exceedance probability of one. **f & g**. Regression analyses to predict the proportion of optimal behaviour (“paired-persist”) from the influence of both the history of the same pairs (experience) and other pairs (inference) for participants’ real data (**f**) and model-simulated data (**g**). Participants relied on both experience and inference to guide their actions, regardless of whether all trials or only the learning period (the first 15 trials of each block) were analysed. Only the Structure Model successfully replicated participants’ behavioural patterns. Grey dots represent the individual regression estimates. The bar plots indicate group mean ± SEM. \**P*<0.05, \*\*\**P*<0.001.

Despite the reversals, participants successfully learned the relational structure between characters, as reflected in their ‘persist or change’ choices. To maximise reward, participants should make initial predictions based on paired transitions, which are of higher probability. Thus, if the expected paired transition indeed occurs, the optimal behaviour is to persist with the initial prediction. However, if a rare transition occurs instead, the optimal behaviour is to change the initial prediction. This is because the task design ensures that characters connected to the same predecessor via different transition types always have opposite preferences (Fig. S2a). For instance, if *A*-to-*B* is a paired transition and *A-*to-*C* is a rare transition, *B* and *C* would always have opposite preferences (Fig. S2a). This design allowed us to determine whether an initial prediction was based on a paired transition— a key indicator of participants’ understanding of the current transition structure. Participants’ behaviours aligned with the optimal behaviour: they persisted significantly more during paired transitions than rare transitions (paired t-test: *t*(35)=10.063, *P=*7.192×10^−12^), and those who adhered more closely to this optimal behaviour achieved higher scores (Pearson’s *r*=0.404, *P=*0.015; Fig. 1f). In addition, participants also successfully learned each character’s preference, as their final predictions reflected the true preferences across blocks (Fig. 1g).

### Relational knowledge supports inference and flexible learning

Having confirmed that participants learned the transitions between characters, we next investigated whether the underlying learning process is facilitated by relational knowledge. To this end, we first assessed how participants learned character transitions through transition history (Fig. 2a). Optimal behaviour reflected participants’ knowledge of character transition structure: if they learned the structure, they would base their initial predictions on paired transitions and persist with them when the expected transitions occurred (‘paired-persist’). Notably, ‘paired-persist’ behaviour relies solely on understanding of the transition structure, whereas ‘rare-change’ behaviour also requires knowledge of character preferences; therefore, we only used ‘paired-persist’ as our measure (Fig. S3a). We tested this by regressing the ‘paired-persist’ behaviour against the number of paired transitions in the recent three trials (LM1, Methods). Indeed, the more paired transitions participants encountered in the recent past, the more likely they were to persist in a current paired transition (*β*+CI=0.222+0.275, *P=*6.19**×**10^−16^; Fig. 2b). However, it is crucial to further distinguish between learning from experience and learning by inference, depending on whether the recent history involves the same pair of characters as the current transition. If the history involves the same pair, learning is experience-based (e.g., learning *A* and *B* are paired through frequent *AB* transitions). If it involves the other pair, learning is inference-based (e.g., inferring that *C* and *D* are paired through frequent *AB* transitions), which requires prior knowledge of the two-pair structure (Fig. 2c).

Can participants actually learn through structure-based inference? For example, even if participants have never directly experienced *CD* transitions, can they leverage their prior relational knowledge to infer that *C* and *D* are paired based on frequent *AB* transitions? If so, seeing *D* follow *C* should not be unexpected, and participants should persist with their initial prediction. Using a separate linear model (LM2, Methods), we found that this is indeed the case: both recent experience of same-pair history (*β*+CI=2.187+0.283, *P*=9.77×10^−15^) and inference from other-pair history significantly increased participants’ likelihood of optimal ‘paired-persist’ behaviour (*β*+CI=2.183+0.381, *P*=1.03×10^−8^; Fig. 2f). The effects held when focusing on the early learning period (first 15 trials) of each block—periods of highest learning demand due to either a fresh start or a reversal— demonstrating that participants rapidly integrated both direct experience (*β*+CI=1.522+0.350, *P*=1.35×10^−5^) and structure-based inference (*β*+CI=1.584+0.624, *P=*0.011; Fig. 2f). These results suggest that people can leverage the relational knowledge to support inference, thereby enabling rapid adaptation to changes throughout the task.

### Reinforcement learning with structure-based inference

To probe the computational mechanism underlying the learning process, we implemented three reinforcement learning (RL) models^24^, each reflecting a distinct learning hypothesis (Fig. 2d; see Methods): The *Frequency Model* assumes that the most frequently observed character in recent trials is also most likely to appear again next, relying solely on the frequency of each character’s occurrence without considering transitions (as a model-free approach; Fig. S4a). The *Transition Model* recognises that the sequence follows a pattern and tracks transition probabilities between characters, but does so naïvely by updating only the transitions it has directly experienced. For instance, observing an *A*-to-*B* transition only strengthens the *A*-*B* edge, without generalising to other edges. The *Structure Model* extends the *Transition Model* by incorporating the two-pair relational knowledge, enabling it to integrate both direct experience and structure-based inference (Fig. 2c). It updates observed transitions bidirectionally (e.g., *A*-to-*B* strengthens both *A*-*B* and *B*-*A* edges) and simultaneously adjusts the unobserved, inferred transition based on the two-pair structure (e.g., *A*-to-*B* also strengthens *C*-*D* and *D*-*C* edges; Fig. 2d). All models implemented learning via the Rescorla–Wagner update rule^25^. The same rule was also applied to the preference learning component of all models, reflecting how participants learn each character’s preference from history (Fig. S4c,d). Model fitting results showed that the *Structure Model* provided the best fit to participants’ behaviour, achieving an exceedance probability of one (Fig. 2e), suggesting that participants used both experience and structure-based inference to facilitate learning as context changed.

Furthermore, regression analyses (LM2) on simulated data showed only the *Structure Model* successfully replicated participants’ behavioural patterns, learning from both direct experience (*β*+CI=1.393+0.119, *P*<10^−16^) and structure-based inference (*β*+CI=1.067+0.167, *P*=1.54×10^−10^; Fig. 2g). In contrast, the *Transition Model*, relying only on the same-pair experience (*β*+CI=0.746+0.103, *P*=5.67×10^−13^), showed no inference from other-pair history (*β*+CI=0.090+0.157, *P=*0.567). The *Frequency Model*, meanwhile, failed to learn even from same-pair history (*β*+CI=-0.007±0.120, *P=*0.953), and other-pair history had a negative impact (*β*+CI=-0.989±0.190, *P*=2.00×10^−7^) due to its incompatible frequency-based assumption (Fig. 2g). These simulation results also held when analyses were restricted to the early learning period (first 15 trials) of each block (Fig. S4a).

Together, these results reveal that, like the *Structure Model*, human learners integrate relational knowledge with direct experience to flexibly infer unseen connections, thereby facilitating learning and adaptation in new contexts. Having established the computational mechanisms of learning by structure-based inference, we then examined how the brain represents this relational structure to implement these computations.

### Relational structure representations in the hippocampus

We next used representational similarity analysis (RSA)^26^ to examine how the brain, particularly the hippocampus (HPC), represents the relational structure. We focused on brain regions identified by whole-brain GLMs—including HPC—that tracked task-relevant information such as character and outcome prediction errors (GLM1-4; Fig. 3a & S4h), as seen in previous studies^27^. We hypothesised that if a brain region represents the relational structure, it would show more similar neural activity patterns for characters connected with higher transition probabilities. To test this prediction, we constructed a trial-level behavioural representational dissimilarity matrix (RDM) for each block, where dissimilarity was defined as one minus the transition probability between characters (e.g., 0.2 for paired transitions, 0.8 for rare transitions, 1.0 for unconnected characters; no self-comparisons; Fig. 3b). We constructed our behavioural and neural RDMs at the trial-level that controlled for the temporal proximity of paired transitions (see Methods, RSA). Neural RDMs were created by measuring distances between trial-level beta estimates (Fig. 3b) across our regions of interest (ROIs; Fig. 3a). We then assessed correlations between the behavioural and neural RDMs to quantify relational structure representations.

**Figure 3.**
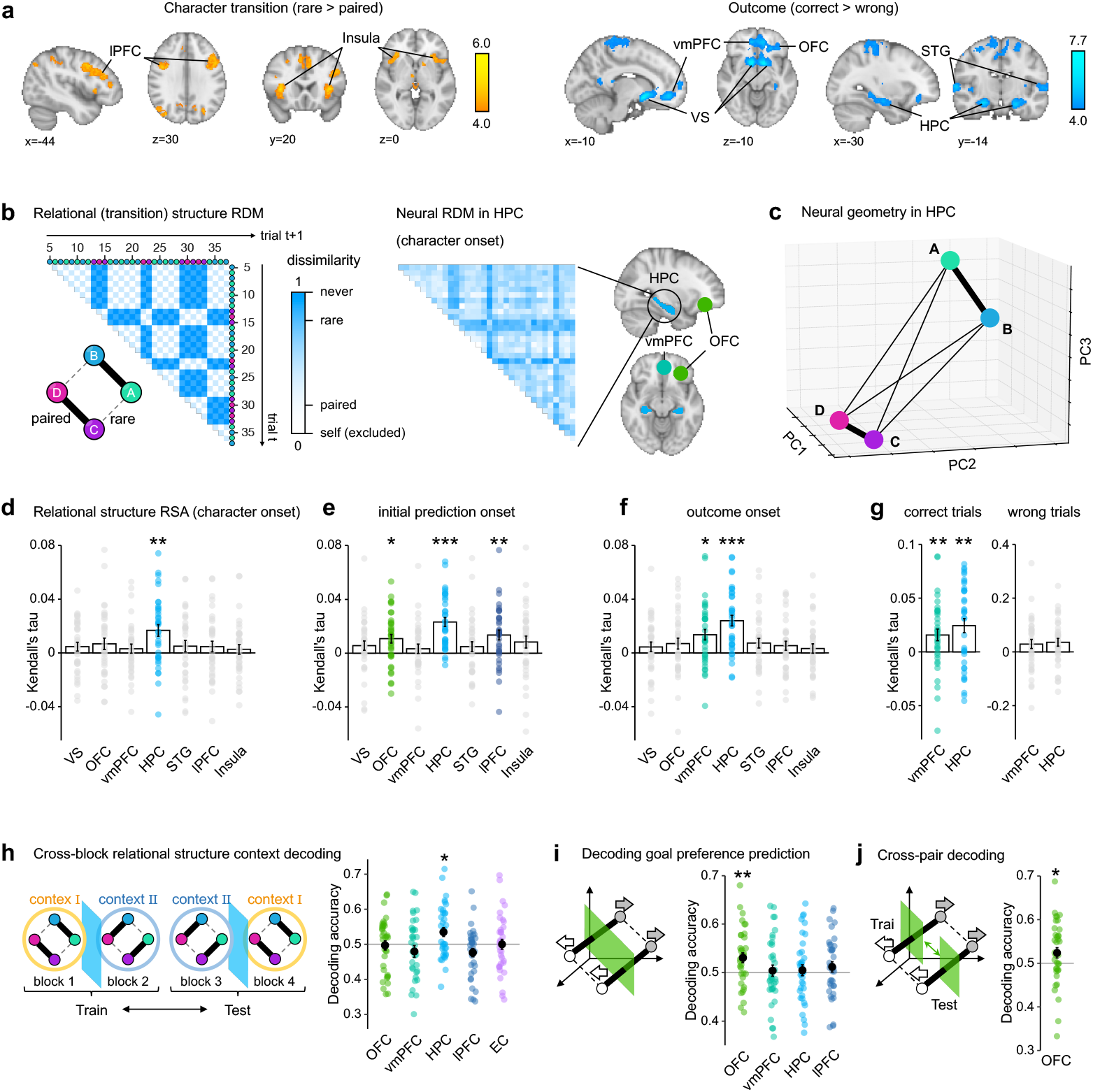
Identification of representation of relational structure in HPC. **a**. Identification of brain regions encoding task-relevant information. Using whole-brain General Linear Models (GLMs), we found activity in prefrontal regions, including an area in and adjacent to the inferior frontal sulcus and lateral prefrontal cortex (referred to as lPFC; left peak Z=5.52, MNI: x=-44, y=8, z=38; right peak Z=5.36, MNI: x=44, y=4, z=30), and insula (left peak Z=5.50, MNI: x=-36, y=20, z=6; right peak Z=5.61, MNI: x=36, y=20, z=4) was stronger during rare transitions than paired transitions. Brain regions including ventral striatum (VS; left peak Z=7.42, MNI: x=-11, y=8, z=-10; right peak Z=7.54, MNI: x=10, y=8, z=-10), ventromedial prefrontal and adjacent medial orbitofrontal cortex (referred to as vmPFC; left peak Z=6.38, MNI: x=-10, y=-46, z=-8; right peak Z=6.32, MNI: x=12, y=48, z=-8), orbitofrontal cortex in and adjacent area to the lateral orbital sulcus (referred to as OFC; left peak Z=5.46, MNI: x=-25, y=35, z=-12), superior temporal gyrus (STG; left peak Z=5.63, MNI: x=-64, y=-18, z=4; right peak Z=6.32, MNI: x=64, y=-16, z=4), and hippocampus (HPC; left peak Z=6.51, MNI: x=-22, y=-15, z=-20; right peak Z=6.64, MNI: x=26, y=-14, z=-18), showed greater activity for correct outcomes than wrong outcomes. See Fig. S4h for model-driven analyses. **b**. Illustration of trial-level RSA. The left panel shows a behavioural RDM, where each entry reflects the dissimilarity between two trials, defined as one minus the transition probability between their respective characters. Self-comparisons (dissimilarity=0) were excluded. The right panel shows the corresponding neural RDM derived from HPC activity, time-locked to character onset. **c**. Example of neural geometry in HPC based on PCA. Each point represents the average neural representation of a character across trials within a single block. Thick edges connect paired characters, which are also represented more closely in the neural space. The model and neural RDMs, along with the neural geometry results, are derived from a single block of data from Subject 1. All example data shown are from a single block of Subject 1, including the model RDM, neural RDM, and PCA results. **d-g**. The relational structure of RSA effects is measured as the rank correlations (Kendall’s tau) between the model and neural RDMs obtained from different ROIs. These correlations were computed for neural activity time-locked to initial prediction onset (**d**), character onset (**e**), and outcome onset (**f**). The HPC consistently represented the relational structure across all phases. During the outcome phase, the RSA effects were further separated into correct (left) and wrong (right) trials (**g**). Significant relational structure representations in HPC and vmPFC emerged only during correct trials. Bars indicate the mean, while error bars represent the standard error. **h**. Cross-block relational structure context decoding was performed by training on half of the blocks and testing on the other half to decode the context (i.e., pairing structure: AB&CD vs. AC&BD) to which each character belonged. ROIs included regions representing relational structures and the entorhinal cortex (EC), which is upstream of the HPC. Only the HPC showed significant decoding accuracy of the relational structure context. **i**. Goal preference prediction decoding was conducted in blocks where paired characters had opposite preferences. Among the selected ROIs, only OFC showed significant decoding accuracy. **j**. Cross-pair decoding in OFC: The classifier was trained to decode goal preference predictions from one character pair and tested on the other pair. OFC still maintained significant above-chance decoding accuracy. Each dot represents an individual. \*\*\**P*<0.001, \*\**P*<0.01, \**P*<0.05.

The activity pattern of the HPC showed a significant correlation with the relational structure at different event points of each trial. At character onset, HPC was the only area to encode the relational structure (Wilcoxon signed rank test *V*=547, *P=*0.003, after Holm-Bonferroni Correction [HBC] for number of ROIs), with no comparable effects in other ROIs (all *P*>0.4, HBC), highlighting its unique role (Fig. 3d and S5b). Our hypothesis-driven identification of HPC representation of relational structure was confirmed by a whole-brain searchlight analysis (Fig. S5a). To visualise such a representation, we conducted a separate principal component analysis (PCA) on HPC activity. The average neural geometry of characters across trials qualitatively mirrored the relational structure in an example block, with paired characters exhibiting more similar patterns (Fig. 3c).

We also examined other time points during the trial. HPC representation was even pronounced during the initial prediction (*V*=646, *P*=7.56×10^−8^, HBC), when participants had to anticipate the next character using the transition knowledge (Fig. 3e). This representation remained significant during the outcome phase (*V*=620, *P*=3.88×10^−6^, HBC; Fig. 3f), indicating a sustained representation across task events. In contrast, some other regions were only engaged during the initial prediction phase, such as the orbitofrontal cortex (OFC: *V*=518, *P=*0.015, HBC) and lateral prefrontal cortex (lPFC: *V*=553, *P=*0.002, HBC; Fig. 3e), suggesting their involvement is limited to the moment when relational knowledge is used for character’s goal preference prediction. The ventromedial prefrontal cortex (vmPFC) was selectively engaged during the outcome phase (*V*=522, *P=*0.014, HBC; Fig. 3f), aligning with its role in encoding both reward occurrence and task structure^3^. Close examination revealed that outcome-phase RSA effects in vmPFC and HPC were observed only in correct trials (vmPFC: *V*=519, *P=*0.003; HPC: *V*=534, *P=*0.002, HBC; Fig. 3g and S6h), not in wrong trials (both *Ps*>0.1, HBC), suggesting that task structure representations might be reactivated by reward delivery^28^ (Fig. S7). All RSA effects remained robust even after reanalysis with partial correlation to control for potential confounds such as predictions and outcomes (Fig. S5c-e). Additionally, we found that the hippocampal RSA effects correlated with various behavioural performances that require knowledge of relational structure (Fig. S5b,f). Together, these results demonstrate the unique and consistent role of HPC in relational structure representation.

### Decoding relational structure context in HPC

Furthermore, we found that HPC not only represented specific character transitions but also encoded the global context in which those characters appeared. The global context is defined by the type of relational structure (*AB&CD* vs. *AC&BD*) to which the current character belonged. Specifically, we trained a classifier to decode the relational structure context using half of the blocks for training and the other half for testing, and found that only the HPC could reliably distinguish between different contexts (*t*(35)=2.779, *P=*0.022, HBC; Fig. 3h and S6). This suggests that even when character identity remained the same, HPC encoded it differently depending on the context in which it appeared. This implies that HPC can track relational structure in a more abstract way that goes beyond specific relations.

### Decoding goal prediction in OFC

However, besides learning relational structure, a second key task requirement is to predict characters’ goal preferences; therefore, we next examined how these goal preference predictions are encoded in the brain. With this aim, we trained a support vector machine (SVM) to decode participants’ final predictions after the character appeared, when participants focused solely on the current character’s goal direction preference. To avoid confounds related to temporal proximity between trials, we restricted our analysis to blocks where paired characters had opposite preferences. Among the ROIs previously identified as representing transitions, only OFC showed significant decoding of goal preference prediction (*t*(35)*=*3.051, *P=*0.008, HBC; Fig. 3i). To test whether such representations in the OFC were independent of specific character pairings, we conducted a cross-pair decoding analysis—training the classifier on one character pair and testing on the other (Fig. 3j; see Methods, Decoding analysis). The OFC again exhibited significant decoding accuracy (*t*(35)=1.956, *P=*0.029), indicating that such representations can generalise across character pairs (Fig. 3j). These findings suggest that OFC encodes character-specific goal preference information in a manner abstracted from the relational structure. While the contributions of HPC and OFC in task representation are often debated^29^, we show that OFC plays a complementary role to HPC: while HPC provides a sustained representation of character-character relational structure, OFC associates each character within this structure with their specific goal preferences and uses it to guide predictions.

### Causal necessity of relational structure representations for inference

Having established that HPC represents relational structures, we asked whether such representations are causally necessary for inference-based learning. To investigate this, we adopted both computational and empirical approaches. Specifically, we used neural network simulations to predict the behavioural effects of disrupting relational structure representations, and then tested these predictions in humans using targeted ultrasound neuromodulation.

### Computational predictions: Lesioning relational structure representation in a neural network disrupts inference

We first asked whether relational structure representations are computationally necessary for flexible learning. To test this, we trained a meta-reinforcement learning (meta-RL) agent on a state-prediction task to model the HPC-like acquisition of relational knowledge across different contexts^29^, mirroring the structure learning in the *Goalkeeper Game* (Fig. 4a). This enabled us to identify specific units representing relational structure in the network and assess whether disrupting them would affect learning. Specifically, the agent was implemented as a recurrent neural network (RNN) with a single hidden layer of 64 long short-term memory (LSTM) units, trained using the advantage actor-critic (A2C) algorithm^30^ (Methods). The environment required the neural network to continuously predict the next state (character) based on the previous state. State transitions were governed by the same two-pair structure as in the *Goalkeeper Game*—a four-state Markov chain containing two paired (80%) and two rare (20%) transitions (Fig. 1c)—sampled from all possible pairings (Fig. 4a), with two unsignaled transition reversals occurring at random time points within each 160-trial episode.

**Figure 4.**
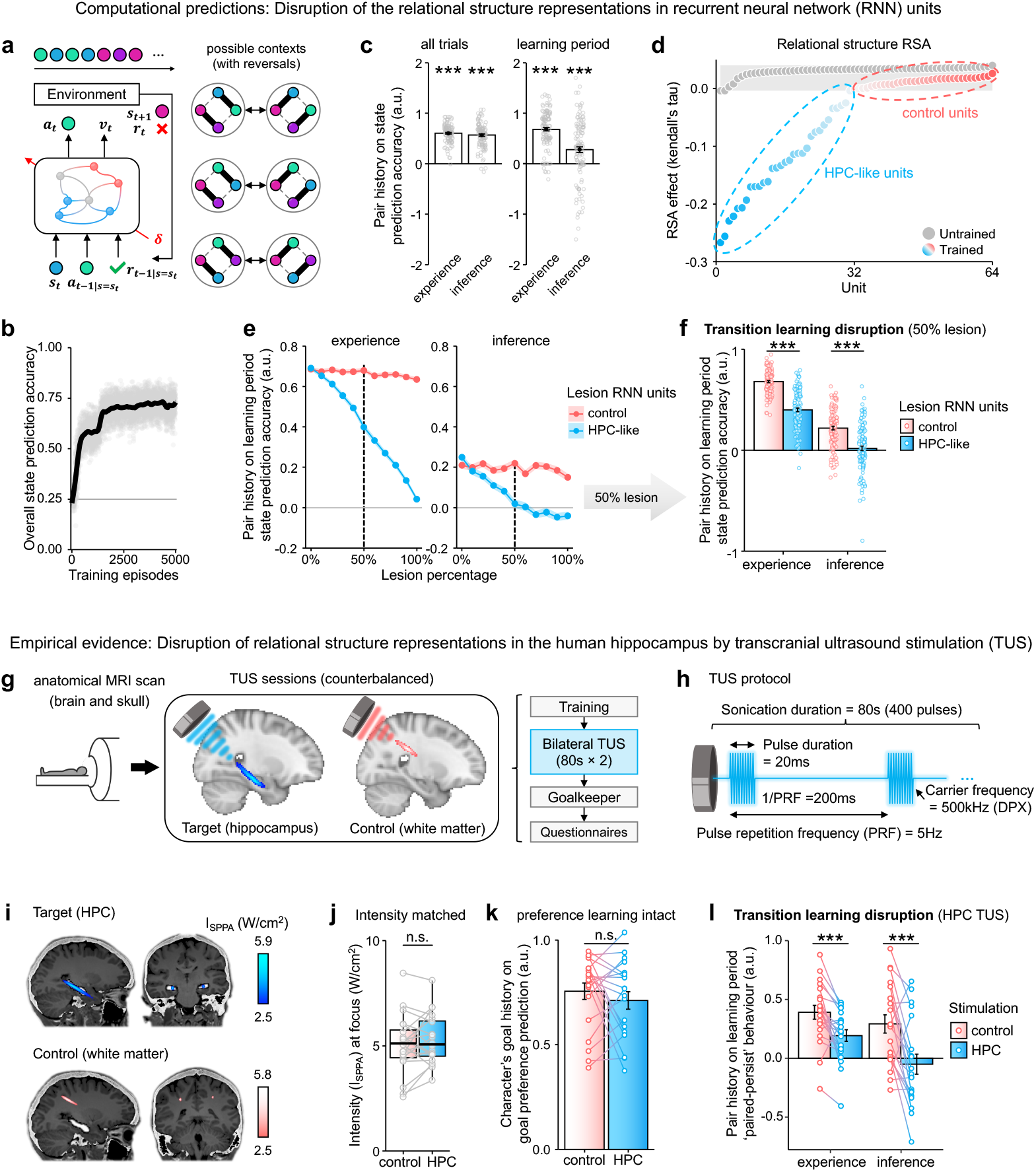
Causal necessity of relational structure representation for inference, in human hippocampus and neural network. **a**. Schematic of the meta-RL neural network architecture: It consisted of a single hidden layer of 64 LSTM units, trained using the advantage actor-critic algorithm. The agent learns to predict the next state, like human participants predicting the next character, by interacting with the environment. At each step, it receives the current state (s_t_), along with the previous action and reward (a_t_−_1_, r_t_−_1_ | s = s_t_) when the same state was last encountered. The agent then generates a current action (a_t_) as a prediction of the next state, and computes a current value estimate (v_t_). The state sequence generated by the environment follows the same two-pair transition structure as the Goalkeeper Game used with human participants. The environment contains all possible pairing contexts (with reversals) and will randomly select one for each episode, mirroring the meta-learning scenario. **b**. The state prediction accuracy increased over training episodes, indicating the network successfully learned the underlying relational structure. Each translucent grey dot shows the accuracy from a single episode; the black line represents the moving average across 500 episodes. **c**. Simulated behavioural data from the trained network. Like human participants, both same-pair (experience) and other-pair (inference) history influenced the network’s prediction accuracy in current paired transitions, regardless of whether the assessment was made across all trials or only during the early learning period. Bars indicate means; error bars denote standard errors. Each dot represents one run of the simulation. **d**. Transition-encoding units identified using trial-level RSA. In the untrained network (grey), the RSA effects of units reflect noise-level fluctuations. In contrast, in the trained network, around half of the units (blue) encoded relational structure, which were referred to as HPC-like units, while the rest (red) were control units. **e**. Lesioning HPC-like units impacts transition learning. As the percentage of randomly lesioned HPC-like units (blue) increases, the network’s use of both experience and inference for predictions during the learning period declines, while lesioning control units (red) has minimal impact. Shaded areas represent standard errors. **f**. A 50% lesioning of HPC-like units (indicated by dashed lines in **e**) significantly reduced the influence of both experience and inference compared to control lesions, with the influence of inference no longer significantly different from zero. \*\*\**P*<0.001. **g**. TUS study procedure and group-level TUS localisation. The TUS study included an anatomical MRI session for simulation-based planning and neuronavigation, followed by two TUS experiment sessions (one week apart). In each session, participants completed the Transition Training, received bilateral TUS targeting either HPC or a white matter control site in a counterbalanced order, performed the 1-h behavioural version of the Goalkeeper Game, and completed post-TUS questionnaires. The averaged intensity map across participants shows the average TUS focus locations in standard space for the target site (HPC, blue, MNI: x=24) and control site (white matter, pink; MNI: x=26). Because individual maximum intensities may not overlap in standard space, the group average map does not necessarily reflect the mean of the participants’ maximum intensities (see **j** instead). The colour indicates scaled group intensity, with the brightest colour representing 1 (highest) and the threshold set at 0.5. **h**. TUS protocol. Participants received 80 s of repetitive stimulation *per* hemisphere, consisting of 400 pulses. Each pulse lasted 20 ms and was repeated every 200 ms, resulting in a pulse repetition frequency (PRF) of 5 Hz. TUS was delivered at a carrier frequency of 500 kHz. **i**. TUS planning of a representative individual. The intensity field map overlaid on brain and skull images (highlighted in white) shows reliable targeting of HPC (blue, upper row) and the white matter control site (pink, lower row). The colour map represents the spatial peak pulse average intensity (I_SPPA_), with a threshold of 2.5 W/cm^2^. **j**. The boxplot displays the average peak intensity (I_SPPA_) at the focus across participants, with dots connected by lines representing individuals. No significant difference in intensity was observed between the two conditions. **k**. Preference learning remained unaffected. The influence of the character’s recent goal direction history on the current goal preference prediction did not significantly differ between HPC and control TUS conditions. **l**. Relational structure learning is disrupted. The influences of experience and inference on participants’ optimal behaviour (i.e., “paired-persist”: persisting in the current paired transition, an index of their relational knowledge) were significantly disrupted by HPC TUS compared to control TUS. Notably, the influence of inference was no longer significantly different from zero following HPC TUS. \*\*\**P*<0.001; n.s., not significant.

At each time step, the network received a one-hot input indicating the current state, the last action–reward outcome in that state, and the trial index, and produced two outputs: an actor head predicting the next state and a critic head estimating the value of the current state (Methods). During training, synaptic weights were fixed within each episode, forcing the network to rely solely on dynamic changes in the network activity, reflecting accumulated experience (fast learning). Weights were updated only across episodes to improve the policy (slow learning). After 5,000 episodes, the network achieved near-optimal performance (∼80% accuracy; Fig. 4b). Like human participants, both experience-based and inference-based influences on the network’s predictions were significant, both across all trials (experience: *β*+CI=0.624+0.018, *P*<10^−16^; inference *β*+CI=0.612+0.031, *P*<10^−16^), and during early learning periods—first 15 trials after a fresh start or a reversal (experience: *β*+CI=0.763+0.041, *P*<10^−16^; inference: *β* ±CI=0.440±0.069, *P*<10^−16^; Fig. 4c).

Given the network’s success in learning the state transitions, we then applied trial-level RSA (see fMRI analyses) to each hidden unit to identify those encoding the underlying relational structure. We again constructed trial-level model RDMs inversely reflecting transition probabilities between states and computed trial-level neural RDMs for each hidden unit (Fig. 3b; Methods). Unlike fMRI multivariate analyses, the RSA was applied separately to each individual unit. We found that around half of the units in a trained network, compared to an untrained network, encoded the relational structure and thus resembled human HPC (Fig. 4d). The rest resembled other brain regions that did not encode the relational structure. But unlike the human HPC, these units encoded the relational structure by exhibiting greater activity changes during paired transitions (Fig. S8a-c), reflecting known disparities in information processing between artificial and biological systems^31,32^.

After identifying HPC-like and control (non-HPC-like) units in the trained network, we separately lesioned each group of units by randomly disabling a subset of them during task performance to test their role in transition predictions. Both experience-based and inference-based influences on predictions during the early learning period gradually declined as the proportion of lesioned HPC-like units increased (simple main effect experience: *F*(1,1316)=656, *P*<10^−16^; inference: *F*(1,1316)=71.78, *P*<10^−16^, HBC), whereas lesioning control units had no comparable effect (experience: *F*(1,1316)=4.544, *P=*0.066; inference: *F*(1,1316)=2.536, *P=*0.112, HBC; Fig. 4e and S8d-f). For example, randomly lesioning 50% of HPC-like units was sufficient to diminish the network’s use of experience (two-sample *t*-test: *t*(198.08)=-13.604, *P*<10^−16^) and inference (*t*(229.73)=-7.222, *P*=7.47×10^−12^), with inference being completely abolished (Bayes Factor compared to zero: *BF=*0.150; Fig. 4f). In contrast, lesioning 100% of HPC-like units abolished both experience and inference (Fig. 4e).

The network simulations confirmed that our trial-level RSA approach can successfully identify neural representations of relational structure that emerge when the structure is learned to support predictions. Critically, disruption of these HPC-like units that represent relational structure impairs both the use of experienced and inferred knowledge for predictions, but especially the latter. These results suggest a clear prediction: if any disruption, even if partial, were applied to human HPC, it would particularly impair structure-based inference during learning. We next examined whether this was the case empirically.

### Empirical evidence: Transcranial ultrasound stimulation of human HPC disrupts structure-based inference

To empirically examine the causal role of the HPC in structure-based inference, we conducted another experiment (Experiment 2) using transcranial ultrasound stimulation (TUS) to modulate its activity. A new group of participants (*N*=20) received bilateral TUS targeting HPC and control white matter sites (hippocampus-unrelated), in two counterbalanced sessions (Fig. 4g). Targeting was guided by individual anatomical MRI scans obtained prior to TUS sessions, using MRI-based neuronavigation and participant-specific TUS simulations, to ensure precise coverage of targeted regions (Fig. 4i). Stimulation consisted of repetitive theta-burst (5 Hz, 20-ms pulses every 200 ms) ultrasound delivered for 80 seconds at a carrier frequency of 500 kHz (Fig. 4h; Methods). The spatial peak pulse average intensity (I_SPPA_) was matched across HPC and control sites, with an average of ∼5 W/cm^2^ (Fig. 4j and S9b). TUS was applied sequentially to the left and right hemispheres (Methods). In each TUS session, participants first completed the *Transition Training* to learn the two-pair relational structure, as in Experiment 1. Then they received TUS on both hemispheres, sequentially, each for 80 s. Immediately after TUS, participants performed the behavioural version of the *Goalkeeper Game* (around 1 hour), followed by post-TUS questionnaires (Fig. 4g).

We found that applying TUS to HPC especially disrupted participants’ learning of character relationships compared to control; the likelihood of persisting during current paired transitions (optimal behaviour) became significantly less sensitive to recent pair history (paired *t*-test: *t*(19)=-2.913, *P=*0.009; Fig. S9d). Specifically, HPC TUS significantly diminished both the experience-based learning (*t*(19)=-3.933, *P*=8.93×10^−4^) and inference-based learning (*t*(19)=-3.999, *P*=7.69×10^−4^), with inference effects no longer distinguishable from zero (Bayes Factor *BF=*0.272; Fig. 4l), indicating a shift from structure-based to simpler, experience-based strategies. Intriguingly, such disruption is best predicted by partial (50%) rather than complete (100%) lesioning of HPC-like units in RNN, suggesting that TUS modulates HPC activity by inducing noise rather than full inactivation^33^. The disruption was most prominent during early learning periods—after fresh starts or reversals—and gradually weakened over time as participants compensated by accumulating knowledge (Fig. S9h). Importantly, HPC TUS had no significant impact on participants’ ability to predict characters’ goal preferences from their recent shot history (*t*(19)=-0.960, *P=*0.350, Bayes Factor *BF=*0.707; Fig. 4k), indicating a selective disruption of structure learning, not preference learning. These findings demonstrate the causal role of HPC in representing relational structure for flexible inference.

We ruled out practice and order effects, as the session order was counterbalanced, and there was no systematic, TUS-independent performance change in experience- or inference-based learning from the first to the second session (Fig. S9g). Overall task performance, including initial and final prediction accuracy, was unaffected by HPC stimulation—likely because the intact goal preference learning compensated for deficits of structure learning. Moreover, transition learning does not always ensure optimal predictions; learners highly sensitive to recent transitions can also be misled by randomness. Given this and the task’s complexity, participants’ initial prediction accuracies were only slightly above chance, making TUS effects difficult to detect (Fig. S9e and S2c). Finally, we conducted a separate behavioural experiment (Experiment 3) to confirm that the observed effects were due to disruption from HPC TUS rather than enhancement from control TUS. A new group of participants (*N*=40) completed the same *Transition Training* and *Goalkeeper Game* in the lab, but without any TUS intervention. Experiment 3 replicated key behavioural findings from Experiment 1 in an independent sample (Fig. S10a,b), supporting the robustness of our results. Most importantly, the influence of experience and inference closely matched those in the control TUS condition of Experiment 2 (Fig. S10c), establishing an independent behavioural baseline and isolating HPC disruption as the cause of structure learning deficits.

## Discussion

Our study shows that humans can quickly learn about relationships in new or changing contexts, not by learning solely from experience, but by inferring from one relationship to another using prior relational knowledge (Fig. 2). Our fMRI findings suggest that such relational knowledge is represented in the hippocampus (HPC). In a simple football-inspired task, HPC persistently represents the relational structure among four characters, despite repeated reversals in the environments (Fig. 3). HPC has long been associated with spatial navigation, and has been suggested to have an analogous function in navigating abstract social and conceptual space^8,9,11,12,34–40^. Intuitively, social navigation shares similarities with spatial navigation. For example, an animal or person can infer a novel route between locations *A* and *C* based on known paths from *A* to *B* and *B* to *C*^7,41^. Similarly, when navigating the social world, we infer relationships between people based on their shared connections (e.g., enemy’s enemy is a friend)^42,43^. Critically, such inference informs learning; when the situation changes and new relations emerge and old ones disappear, people learn quickly by combining what they directly experience with what they infer.

Moreover, as a major advance, our study provides causal evidence that relational structure representations in HPC are necessary for the inference of relationships. Directly testing the causal contribution of HPC in higher cognition has long been difficult in humans. Lying deep in the brain, human HPC has been inaccessible to conventional non-invasive stimulation tools, such as transcranial magnetic stimulation (TMS) or transcranial direct current stimulation (tDCS)^13^. Other indirect approaches either struggle to isolate the HPC’s specific contribution from connected regions or produce modest behavioural effects^44,45^. In this study, we used a new non-invasive method, focused transcranial ultrasound stimulation (TUS), to directly target human HPC with millimetre precision and temporarily modulate its activity. Previous work on animal models has demonstrated the safety, reliability, reversibility, and feasibility of TUS to modulate deep brain regions without affecting overlying areas^14–21^, suggesting its potential for studying cognitive functions in the human brain^46^. By developing a TUS protocol with optimised focal coverage targeting bilateral human HPC (by stimulating along the long axis of the HPC from the back of the head), we disrupted participants’ ability to learn new relationships: while learning from direct experience was partially impaired, inference from prior relational knowledge was entirely abolished (Fig. 4l). These findings provide causal evidence that HPC is essential for flexible, inference-based learning.

Importantly, this empirical study of the causal role of relational structure representations in inference is guided by our predictions from neural network modelling. Theoretical work has suggested that abstract representations emerge in HPC simply through learning to predict the future^47–49^. Our study demonstrates that abstract representations of relational structures emerged in human HPC through trial-by-trial predictions about the next character (Fig. 3). To model this process, we trained a recurrent neural network (RNN) on a similar task to predict the next state through a meta-reinforcement learning (meta-RL) process^29,30^. After training, the RNN developed units that encoded abstract relational structure between states, like human HPC (Fig. 4d). This pattern was observed consistently across networks with varying training parameters (Fig. S8), despite the known disparities between artificial and biological systems in their ways of representing information^31,32^. Like human participants, the RNN also managed to leverage relational knowledge for inference, despite never being explicitly taught the underlying task structure (Fig. 2f and 4c). Lesioning these HPC-like units in the RNN primarily disrupted structure-based inference, mirroring the hippocampal TUS effects (Fig. 4f and 4l), and offering a powerful framework for testing the causal role of neural representations and predicting behavioural effects of targeted disruption^50^. Together, both computational predictions from RNN modelling and empirical results from TUS manipulation demonstrated that representations of relational structure in HPC are causally necessary for inference.

Previous studies on patients with hippocampal lesions have established HPC’s causal role in long-term memory^51^. However, our study expands this perspective by providing direct, non-invasive causal evidence from healthy humans that HPC also represents the structure of the world to support inference. These findings are consistent with the framing of memory as an inferential process: rather than storing all facts exhaustively and in isolation, the HPC abstracts structures from past experiences to predict new experiences^52^. Such representations in HPC are suggested to be abstract and stripped of extraneous details, a property that enables inference and generalisation^53,54^. From this perspective, the memory impairments observed in patients with HPC damage may reflect compromised inference, which can also explain why such patients exhibit difficulties with imagination and planning^55,56^.

Our findings also resonate with recent evidence that HPC supports hidden state inference. In rodents, hippocampal activity reflects inference of latent structures that are not directly observable from sensory input, dynamically remapping representations when these latent structures shift^57,58^. In our task, participants also inferred which of the possible latent pairing structures (*AB&CD, AC&BD*, or *AD&BC*) governed character transitions (Fig. S4e). As these structures varied across blocks and were never directly observable, successful performance depended on inferring them from experience. This ability to identify and update the latent structure, rather than simply learning observable transitions, was central to our *Structure Model*, which best fit participants’ behaviours (Fig. 2 and S4e). Supporting this, decoding analyses revealed that HPC distinguished which pairing structure each character belonged to, reflecting context-dependent coding (Fig. 3h). Together, these results indicate that the HPC not only encodes observable transitions but also infers latent structures, thereby enabling rapid adaptation to hidden changes in the task.

While HPC represented relational structure, two other important task variables were represented in the frontal cortex. First, prediction errors of the next character were associated with increased activity in the left prefrontal cortex (lPFC), consistent with previous studies that have shown its role in representing state prediction errors (Fig. S4h)^27^. Additionally, the orbitofrontal cortex (OFC) predicted each character’s goal preferences (Fig. 3i,j and S5g). Previous studies have implicated OFC in predicting outcomes and supporting goal-directed behaviours^3,59–61^. Although OFC briefly represented the relational structure at the time of initial prediction (Fig. 3e), it shifted to only encode goal prediction once the character appeared. This is because after the character had appeared, the character-character relational structure was no longer relevant, and participants could focus solely on the character’s preference. Such dynamics are reminiscent of OFC’s role in representing stimulus value in a state-dependent manner^62^. By contrast, HPC uniquely and consistently represented relational structures, and such representations remained robust even after controlling for goal preference-related variance (Fig. S5c-f). Together, in line with rodent studies^63^, these results suggest the complementary roles of HPC and OFC: the HPC constructs the fundamental relational structure, while OFC anchors this structure to specific goals and uses it to guide predictions (Fig. 5). Building on this, future studies might apply TUS to OFC to test this complementary role, though its anatomical location near the eyes demands specialised protocols for safe and effective targeting.

**Figure 5.**
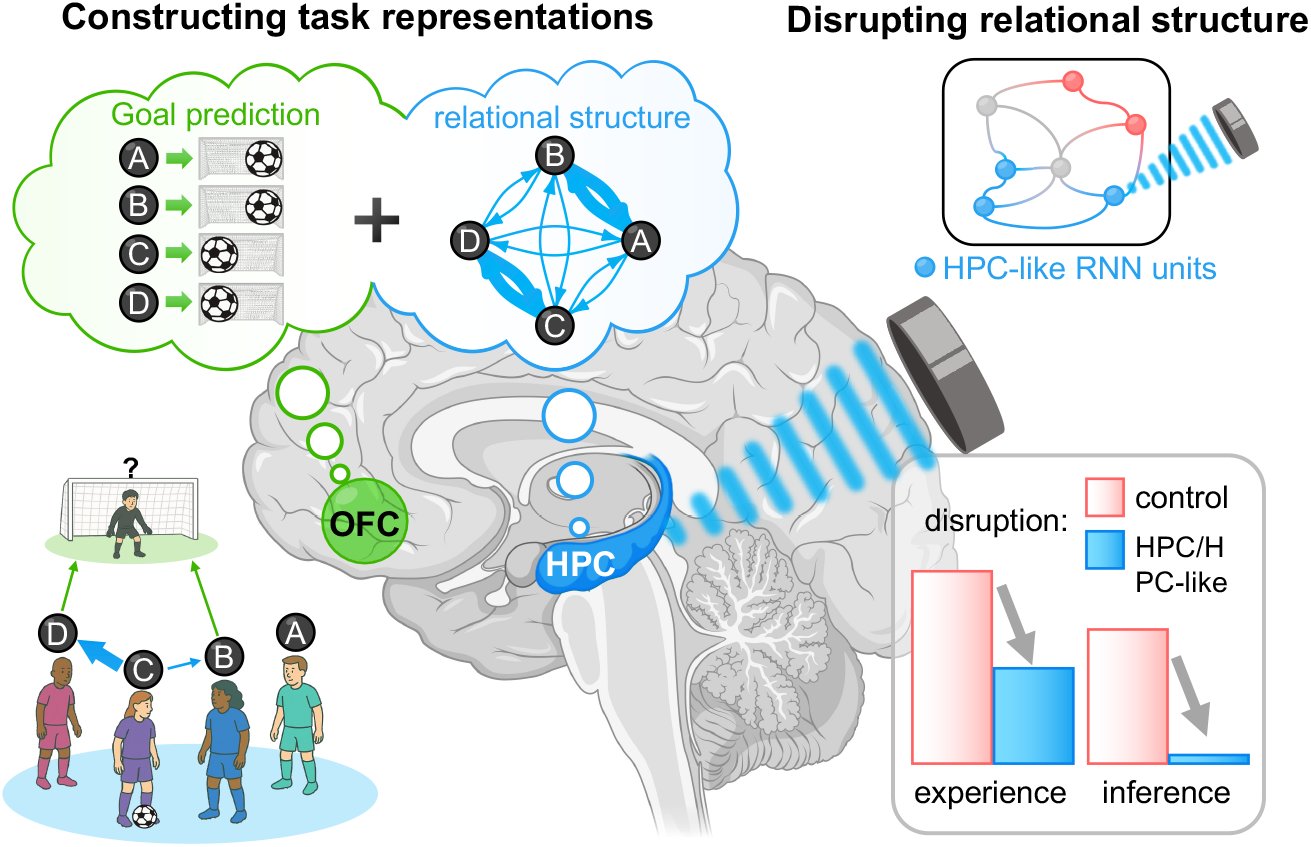
Summary. HPC represents relational structure between characters across contexts, and OFC predicts individual goal preferences. Causal manipulations, including TUS targeting HPC in humans and lesioning HPC-like units in RNNs, disrupt representations of relational structure and impair structure-based inference.

In summary, using a behavioural task combined with fMRI, we demonstrated that HPC represents the relational structure between individuals, which supports inference and facilitates learning in a changing environment. By lesioning the ‘HPC-like’ units in an RNN and using TUS to disrupt activity in human HPC, we demonstrated the causal necessity of relational structure representations in HPC for inference (Fig. 5). More broadly, TUS offers a powerful tool to understand the function of deep brain structures, holding promise for both basic cognitive neuroscience research and future clinical translation.

## Methods

### Overview

We conducted three experiments. The first experiment examined behavioural and neural evidence from functional magnetic resonance imaging (fMRI) for the existence of representation of relational structure, while the second experiment investigated the causal role of the hippocampus in representation of relational structure, by applying transcranial ultrasound stimulation (TUS) to disrupt its activity. A final, third experiment collected additional behavioural data. The three experiments used different samples of participants.

### Participants

#### Experiment 1 (fMRI)

A total of 39 healthy volunteers were initially recruited. Three participants were excluded from the analysis due to excessive head motions (*n*=1) or their performance at chance level (negative scores substantially below zero, *n*=2, one of whom fell asleep in the scanner). The final sample consisted of 36 participants (18 female) aged between 18 and 39 years (age mean ± SD = 24.9 ± 4.6). Participants were paid £60 for their participation in the study, plus a bonus of up to £10, which was calculated based on their performance.

#### Experiment 2 (TUS)

A total of 32 participants initially underwent MRI scans in preparation for TUS sessions. Following K-Plan individualised TUS simulations, only 23 of these participants met the inclusion criteria and were invited to the subsequent TUS sessions. The inclusion requires the spatial peak pulse average intensity (I_SPPA_) for the participant’s target sites greater than 3 W/cm^2^ bilaterally and satisfying all safety criteria (See TUS simulations). Three of these 23 invited participants dropped out after the first TUS session, leaving a final sample of 20 valid participants (11 female) aged between 19 and 33 years (age mean ± SD = 25.4 ± 4.6), who completed both experimental sessions. Participants were compensated £105 for participation (£15/hour for 7 hours), a £50 bonus for completing both TUS sessions, and up to £10 bonus based on performance per session. All participants were pre-screened for contraindications to MRI and TUS. We estimated the required sample size using G*Power^64^, based on a previous TUS study with a similar within-subjects design^65^, and an observed effect size of Cohen’s *d* = 0.77. For a paired *t*-test with 80% power, the minimum sample size was 14 participants at *α*=0.05 and 20 participants at a more conservative *α*=0.01. To ensure the reliability of results, we recruited 20 participants who met the criteria.

#### Experiment 3 (behavioural)

A total of 43 participants initially participated. Three participants who showed chance-level performance (negative scores substantially below zero) were excluded. The final sample included 40 participants (25 female) aged between 18 and 34 (age mean±SD = 23.7±4.1). Participants were compensated £15 for their one-hour participation, and up to a £10 bonus based on performance.

Participants in all experiments were healthy, right-handed, and had normal or corrected-to-normal vision. They had no history or current diagnosis of neurological or psychiatric disorders and were free of psychoactive medications at the time of the study.

#### Ethical compliance

All studies were approved by the Central University Research Ethics Committee (CUREC) of the University of Oxford (Exp.1 & 3: R85346/RE001; Exp. 2: R90784/RE002). Informed consent was obtained from all participants before the experiment.

### Experimental Procedure

#### Experiment 1 (fMRI)

Participants underwent a single fMRI session, during which they completed the fMRI version of the ‘*Goalkeeper Game*’ inside a 3T MRI scanner. The entire MRI scan, including a structural MRI scan and a task-based fMRI scan, lasted approximately 70 minutes. Before the scan, participants completed the ‘*Transition Training*’ and received instructions and practice on how to perform the *Goalkeeper Game*.

#### Experiment 2 (TUS)

The TUS experiment consisted of three sessions: a structural MRI session (30 minutes) and two TUS sessions (each 2.5-3 hours). The structural and zero echo-time (ZTE, or Siemens Petra) MRI scan for each participant was acquired in the first session. This was subsequently used to guide the TUS K-Plan simulation and actual TUS stimulation with the aid of Brainsight neuronavigation in the subsequent two sessions. In the two TUS sessions, participants received TUS, in one case targeting the hippocampus and in the other targeting white matter as a control condition. In each session, participants first completed a *Transition Training*. They next learned how to play the *Goalkeeper Game* through instruction and practice. Then, bilateral TUS stimulations were applied at the designated targets for 80 seconds (offline). After stimulation, they performed the behavioural version of the *Goalkeeper Game* and completed post-TUS questionnaires. The two TUS sessions were scheduled at least one week apart, with the stimulation target (hippocampus, control) order counterbalanced across participants.

#### Experiment 3 (Behavioural)

This experiment consisted of a single session in which participants completed the *Transition Training* followed by the behavioural version of the *Goalkeeper Game* in a testing room.

### Experimental tasks

#### Relational (transition) structure

Before the main task, participants completed three *Transition Training* blocks. Each training block featured a unique set of characters, all distinct from those used in the main *Goalkeeper Game*. The purpose of the training was to teach participants the transition-based relational structure (i.e., transition structure) between characters, which would be beneficial in the subsequent *Goalkeeper Game*. This is because the characters’ transitions in the *Transition Training* and *Goalkeeper Game* shared the same *structure* but used different character sets.

The structure was designed as a four-state cyclical Markov chain with no self-repetition, where each character corresponds to a distinct state (Fig. 1c). The transitions between states (characters) were symmetric and bidirectional, meaning that if a transition from one state to another occurred with a certain probability, then the reverse transition had the same probability of occurrence (if character *A* passed to character *B* with probability, then *B* passed to *A* with probability too). In this structure, the four states were systematically arranged into two pairs. Members of each pair were more likely to follow one another in the sequence, with a transition probability of 0.8 (paired) bidirectionally. Conversely, transitions between unpaired states were either rare (0.2) or non-existent (0). Additionally, no state could transition to itself, ensuring no self-repetition. Notably, the structure was never explicitly shown to participants. Instead, participants had to learn it through trial-and-error observations of transitions between the four characters. During the *Transition Training*, participants were informed that four characters were passing the ball to each other, and the order of ball passes followed a certain “passing pattern.” Participants were informed that despite changes in the specific characters, the underlying “passing pattern” remained consistent across blocks and would also apply in the subsequent *Goalkeeper Game*. Therefore, learning this pattern was beneficial.

#### Transition Training

Each *Transition Training* block consisted of two parts: the *Transition Learning* part and the *Transition Testing* part (Fig. 1d).

In the *Transition Learning* part, participants observed four characters appearing one after another on the screen. On each trial, a character was displayed on the screen for 1 s, followed by a 500 ms inter-trial interval. The next character was probabilistically selected based on the predefined transition probabilities: 0.8 if it was paired with the current character, 0.2 if it was an unpaired but connected character, and 0 if there was no connection. To ensure participants were engaged, the football icon appeared above the character’s image on 50% of randomly selected trials, prompting participants to press a response key as quickly as possible. Each *Transition Learning* sequence consisted of 60 trials, immediately followed by *Transition Testing*.

In the *Transition Testing* part, we assessed participants’ understanding of the “passing pattern” (i.e., transition structure). To this end, participants were presented with a character and asked, “Who is more likely to follow this character to receive a pass?” Participants chose between two other characters. Consider a scenario with four characters (*W, X, Y, Z*) where *WX* and *YZ* are paired transitions (0.8 probability), and *W-Y* and *X-Z* are rare transitions (0.2 probability; Fig. 1d and S1a). In this case, if the current character is *X*, the likelihood of transitioning to the next character follows the order: *W* (0.8) > *Y* (0.2) > *Z* (0). Thus, if participants were tested, ‘Who is more likely to follow *X* to receive a pass?’ and given a choice between *W* and *Y*, the correct answer should have been *W*; when choosing between *Y* and *Z*, the correct answer should have been *Y*. Participants were asked eight questions, and no feedback was provided after they responded. They only proceeded to the next block if they achieved 90% accuracy (a minimum of 7 correct answers). Otherwise, they had to repeat the whole block. Participants were allowed as many attempts as they needed to pass all three training blocks before proceeding to the main task.

In Experiments 1 and 3, participants completed three *Transition Training* blocks once before the main task. In Experiment 2, participants completed three *Transition Training* blocks before the main task in each experimental session (visit).

### Goalkeeper Game (main task)

#### Overall description

Having learned the general two-pair transition structure between the characters, participants started the *Goalkeeper Game*. In the *Goalkeeper Game*, the participants played the role of a goalkeeper, while four new characters (*A, B, C*, and *D*) were introduced as shooters, taking turns to shoot the ball. Participants were instructed to learn the “passing pattern” (transition structure) between characters, which followed the same structure as in the *Transition Training*. Additionally, each character had a preferred goal direction (left or right), which participants had to learn during the experiment, along with the transitions between characters. Participants were told this was akin to a free-kick scenario, where goalkeepers had to predict the direction before knowing which character would take the shot. To make accurate predictions, they had to rely on two pieces of information they could acquire during the task: (1) to internally anticipate the likely next character in trial *t*, using the identity of the previous character (in trial *t*-1) based on the learned two-pair structure, and (2) applying their knowledge of the anticipated character’s goal preference to guide their predictions about the direction. The trial timeline is detailed below (Fig. 1a).

#### Trial timeline

At the start of each trial, a silhouette of the goalkeeper was displayed on the screen, followed by the appearance of a question mark and two arrows indicating left and right directions for 0.5 s. Upon seeing the arrows, participants then had 3 s to make an initial prediction by pressing a button to indicate whether the ball would be shot to the left or right. As participants did not know the identity of the character who was shooting, they had to make an initial prediction by anticipating the next character based on the transition structure. Having anticipated the next character, they could predict the goal direction from their knowledge of that character’s goal preference. Each character preferred a goal direction (left or right) for shooting with 0.8 probability and the other direction with probability 0.2. After participants made their initial prediction, their chosen direction was highlighted in grey and displayed for a jittered inter-stimulus interval (ISI) of 3 to 7 s (average: 4.8 s), generated using a truncated exponential distribution (*μ* = 6). After the delay, the photo of the character appeared on the screen for 1.5 s. During this time, participants were allowed to revise their initial prediction: having observed the character, they could either persist with or change their initial prediction. Again, they could indicate their predicted direction by pressing the left and right arrows for left and right directions, respectively. If no response was made within the time limit, participants proceeded to the next phase of the trial, and their initial prediction was automatically recorded as their final prediction. For example, if a participant initially predicted left, they could change to the right by pressing the corresponding button or persist with their initial prediction by pressing the same button or taking no action. Once the final prediction was confirmed, it was displayed in black. After another jittered ISI (3-7 s, average: 4.8 s, truncated exponential distribution with *μ*=6), the outcome (the actual goal direction of the character) was shown on the screen for 2 s. A football icon on the left or right side of a goal indicated the direction of the character’s shot. In addition, an arrow appeared on the predicted goal direction whose colour signalled how many points participants collected on that trial: green (+2), yellow (0), and red (−2). The payoff matrix is detailed below. After a jittered inter-trial interval (ITI) of 3.5 to 5.5 s (average 4.8 s), generated using a truncated exponential distribution (*μ* = 4), the next trial began. In the behavioural version of the *Goalkeeper Game*, which was used in Experiments 2 and 3 without fMRI scan, we removed the jitters that were necessary for fMRI. We fixed the ISIs and ITIs to 1 s and 2 s, respectively, and participants were given 5 s for the initial prediction and 2 s for the final prediction.

#### Payoff matrix

Participants were told that persisting with their decision was like making a “diving save” in a football game. In this case, a correct prediction that matched the character’s actual goal direction resulted in a perfect diving save with a gain of two points (+2, green arrow), while a wrong prediction incurred a loss of two points (−2, red arrow). If participants changed their prediction, they were informed that they would make a “slow save.” Here, a correct prediction (initially wrong but changed to correct) resulted in a slow but safe save with no point change (0 points), but a wrong prediction still led to a loss of two points (−2, red arrow). Participants were encouraged to collect as many points as possible.

In some cases, if participants failed to respond within the time limit for the initial prediction, they were still required to make a final prediction during the subsequent character epoch. If they managed to do so, the final prediction was treated as a change of their initial prediction (meaning that a correct final prediction led to a 0-point, and a wrong prediction led to a 2-point loss). However, failing to make a final prediction again after missing the initial prediction resulted in the participant making no save and incurring a loss of 4 points. This more severe punishment served as a reminder for participants to stay focused on the task. Participants were pre-emptively informed that the experiment would be terminated if they made no saves more than three times.

#### Task design & schedule

The *Goalkeeper Game* contained 160 trials, divided into four blocks of approximately 40 trials each. Character transition structure and characters’ goal preferences underwent unsignaled reversals. There were two “transition reversals” and one “preference reversal” over the course of the task (Fig. 1e).

For “transition reversals,” in each block, the sequence of four characters followed the two-pair transition structure (see ‘Relational (transition) structure’ above for details): they form two pairs, with 0.8 transition probability between two paired characters, and transitions between unpaired characters were either rare (0.2) or non-existent (0). Importantly, however, two transition reversals occurred at the start of blocks 2 and 4, resulting in an alteration in the transition probabilities: previously paired transitions (0.8 probability) became rare transitions (0.2 probability), and vice versa (i.e., previously rare transitions became paired transitions). These reversals maintained the overall structure by ensuring that there were always two pairs. As a result, blocks 2 and 3 shared the same transition structure, as did blocks 1 and 4. Participants were not explicitly informed about the structure of the sequence or the reversals of structure across blocks. They were only instructed that the characters’ sequence, or their “passing pattern”, followed a rough order, similar to the one they had experienced in the initial *Transition Training* blocks. They also received guidance that the rough order could be learnt by observation but might change occasionally.

One “preference reversal” occurred between blocks 2 and 3, where the preferred goal directions of two characters were reversed (from left to right or *vice versa*). As mentioned above, each character preferred a direction (left or right) with 0.8 probability while shooting in the other direction with probability 0.2. Throughout the experiment, two characters, *A* and *D*, maintained their goal preferences, with A preferring the right and *D* preferring the left. In contrast, characters *B* and *C* swapped their preferences during the preference reversal: *B*, who initially preferred left, switched to the right, and *C* changed from right to left. Consequently, in each block, there were always two characters preferring the right and two preferring the left.

In the fMRI version of the *Goalkeeper Game*, we generated 16 distinct character sequences, each with corresponding goal direction outcomes based on the above design, and randomly assigned each participant to one of these sequences. Notably, half of the participants were assigned to sequences starting with *AB & CD* pairs, while the others were assigned to sequences starting with *AC & BD* pairs. In the first block with *AB & CD* pairs, paired characters have opposite goal preferences (e.g., *A* and *B* are paired, with *A* preferring the right and *B* preferring the left). In the first block with *AC & BD* pairs, paired characters have the same goal preference (e.g., *A* and *C* are paired, both preferring right). Therefore, although all participants were tested with the same general task design, it was instantiated in slightly different ways. In the behavioural version of the *Goalkeeper Game* (Experiments 2 and 3), participants played two distinct sequences per session, each featuring a different character set, assigned pseudo-randomly. In Experiment 2 (TUS), participants completed two sessions and thus played the game four times in total, with each game using a unique sequence and character set to prevent repetition. Furthermore, to minimise any potential impact of the facial features of any character, we randomly assigned a set of pre-selected photos to each *Goalkeeper Game* and *Transition Training*. Within each set, the four photos were then shuffled to represent different characters. The face stimuli were obtained from the Chicago Face Database (https://www.chicagofaces.org/) and selected while balancing race and gender^66^.

### MRI data acquisition

#### Experiment 1

MRI data were acquired using a Siemens 3-Tesla Magnetom Prisma scanner with a 32-channel head coil. Slices were acquired in interleaved order with an oblique angle of -30° from anterior to posterior commissure (AC-PC) to reduce frontal signal dropout. BOLD functional images were collected using a multiband gradient-echo EPI sequence with the following parameters: echo time (TE) = 30 ms; repetition time (TR) = 1200 ms; flip angle = 60°; multiband acceleration factor = 3; in-plane (iPAT) acceleration factor = 2; FOV =216mm; voxel size = 2.4 × 2.4 × 2.4mm. Field map images were acquired to reduce geometric distortion using a dual-echo gradient echo sequence with the following parameters: TE1/TE2 = 4.92/7.38 ms; TR = 482 ms; flip angle = 46°; FOV = 216mm; voxel size = 2.0 × 2.0 × 2.0 mm. Finally, T1-weighted structural images were acquired using an MPRAGE sequence with the following parameters: TE = 3.97ms; TR = 1900 ms; Inversion time (TI) = 904 ms; 192 sagittal slices; FOV = 192 mm; voxel size = 1.0 × 1.0× 1.0 mm.

#### Experiment 2

MRI data were acquired for individualised TUS simulation using the same scanner and head coil as Experiment 1. First, T1-weighted structural images were acquired using an MPRAGE sequence with the following parameters: TE = 2.26ms; TR = 2300 ms; TI = 900 ms; 192 sagittal slices; FOV = 256 mm; voxel size = 1.0 × 1.0× 1.0 mm. Then, zero-TE (ZTE) images were acquired using a PETRA sequence with the following parameters: TE = 0.07ms; TR = 3.61ms; TI = 900 ms; 320 transversal slices; slice thickness = 0.75 mm; FOV = 240 mm; voxel size = 0.8 × 0.8× 0.8 mm^67^.

### Transcranial ultrasound stimulation (TUS)

#### TUS localisation

The initial target coordinates, defined in Montreal Neurological Institute (MNI) space, were: left hippocampus (HPC): x = -28, y = -20, z = -18; right HPC: x = 26, y = -18, z = -20; left white matter (WM): x = -27, y = -32, z = 28; right WM: x = 24, y = -33, z = 29. The white matter we chose was a tract unrelated to the hippocampus. Target locations were optimised in each participant’s native space based on their T1-weighted MRI data and refined with Brainsight neuro-navigation software v2.5.3 (Rogue Research Inc.). Transducer placement was determined by minimising the angle between the transducer exit plane and the skull surface, thereby reducing acoustic reflection and improving transmission efficiency (Fig. S9a). Additionally, positioning aimed to align the ultrasound beam along the elongated axis of the hippocampus to optimise focal coverage. The transducer-to-skull coupling gap was considered during the calculation of transducer coordinates. Multiple potential transducer locations were evaluated, and the final selection was based on simulation results.

#### TUS protocol

TUS was delivered using the Neurofus PRO TPO system with the four-element annular DPX-500-4CH transducer (Sonic Concepts, Brainbox Ltd., Cardiff, United Kingdom; carrier frequency: 500 kHz; diameter: 64mm). The DPX-500-4CH transducer is designed to target deep brain sites with a maximum focal depth of 125.00 mm. The Neurofus PRO TPO system driving the DPX transducer was calibrated prior to the study, using hydrophone scanning in a water tank^68^. We used a theta-burst patterned TUS protocol with the following parameters: pulse duration = 20 ms, pulse repetition interval = 200 ms, duty cycle = 10%, and total duration = 80 s, giving a total of 400 pulses (Fig. 4h). For hippocampal stimulation, the free-field spatial peak pulse-average intensity (I_SPPA_) was set at 75 W/cm^2^ to compensate for transcranial attenuation. Energy loss occurs due to absorption, reflection, refraction, and scattering at the skull interface, which is unavoidable despite optimal transducer placement. The attenuation rate was around 90%, with final values varying across participants. To ensure matched intensity at the target focus, we individualised the white matter control stimulation by adjusting the free-field I_SPPA_ between 55 W/cm^2^ and 75 W/cm^2^, based on each participant’s simulation results. The focal depth was adjusted for each participant and brain region, respectively.

#### TUS simulations

Simulations were conducted for all participants using k-Plan v1.0.1 (Brainbox Ltd., Cardiff, United Kingdom) to ensure precise targeting and safety. K-Plan simulations primarily relied on pseudo-computed tomography (PCT) images of each participant’s individual skull, converted from PETRA ZTE images using an open-source MATLAB toolbox (https://github.com/ucl-bug/petra-to-ct). All the images and coordinates were transformed to ZTE space for K-Plan simulations using FMRIB Software Library’s (FSL) linear transformations (FLIRT) before simulation^69,70^. We conducted acoustic and thermal simulations using the coordinates and protocol parameters mentioned above. The thermal effects were simulated using K-Plan for the 80-second stimulation period plus a 100-second cooling period after it. All the simulations shown in the results were done before the actual stimulation session. Simulations confirmed the precise bilateral targeting of both the HPC and WM with *in situ* intensities of approximately 5 W/cm^2^ on average (Fig. 4j). Participants with simulated maximum I_SPPA_ at the HPC below 3 W/cm^2^ were not invited to the actual TUS session because we estimated that there might be insufficient acoustic transmission to cause any measurable behaviour change^20^. The mechanical index (MI) remained below 1.9, and the thermal dose did not exceed 0.25 CEM43 (Fig. S9b), ensuring compliance with the safety limits of the International Consortium for Transcranial Ultrasonic Stimulation Safety and Standards (ITRUSST) guidelines^71^.

#### Applying TUS

Before stimulation, each participant’s head was prepared by parting hair over the target regions and applying ultrasound transmission gel (Aquasonic 100, Parker Laboratories Inc.)^68^. A plastic coupling system was attached to the transducer, filled with degassed water, and sealed with a thin latex membrane to ensure effective acoustic coupling. The transducer output, including the water and membrane setup, was pre-calibrated using a hydrophone setup in a water tank to ensure consistent acoustic properties. The membrane was slightly overfilled to compensate for skull curvature and minimise air gaps (Fig. S9a). Neuro-navigation was performed using Brainsight software on each participant’s T1-weighted MRI scan, aligning the transducer precisely to the predefined coordinates. Each TUS session consisted of bilateral stimulation, beginning with the left hemisphere, followed by the right hemisphere, with only a brief interval for localisation adjustments. During stimulation, the transducer’s position was continuously monitored and adjusted in real time. Deviations from the intended coordinates were recorded, with real-time manual corrections ensuring alignment errors remained below 1 mm in position and 1 degree in angle, maintaining high targeting precision. Participants received stimulation at a single brain site per session, with the order counterbalanced across the group. Participants were not informed of the brain regions under stimulation.

#### Post-TUS questionnaire

To assess the potential adverse effects of TUS, participants completed a post-TUS questionnaire immediately after each session and again 24 hours later. The questionnaire evaluated sensory, physical, emotional, and cognitive responses related to the stimulation. Participants rated the intensity of physical symptoms (e.g., headache, tingling, dizziness, nausea, muscle discomfort) and cognitive or emotional changes (e.g., attentional difficulties, mood fluctuations) using a four-point scale: Absent, Mild, Moderate, or Severe. Additionally, they indicated whether symptoms were likely related to TUS (Unrelated, Unlikely, Possible, Probable, or Definite). The questionnaire also included open-ended questions, such as “What did the ultrasound stimulation feel like?” for participants to report any additional experiences or sensations, ensuring a comprehensive and unbiased assessment of TUS effects (Fig. S9c).

### Statistical analysis

We analysed the data using R (R Development Core Team, 2008). We carried out multilevel regression models using the lme4 (version 1.1.27.1) package in R. In each multilevel regression model, we focused on the group-level fixed effects and considered participants as random effects to estimate statistical significance (linear mixed model). For data visualisation (where single-subject data was depicted) and within-subject comparisons, we conducted logistic regression for each participant and plotted each participant’s data. Notably, all the regression results stay consistent regardless of whether by linear mixed regression or by performing regressions for each participant and testing their group-level effects. Here are the linear models:

#### LM1

To investigate how participants learned the transition structure based on paired-transition history, we first used a linear model as follows:

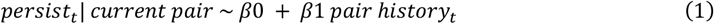

Where “*persist*_*t*_” indicates, on trial *t*, whether participants persisted with their initial goal preference prediction during a paired transition (persisted = 1, changed = 0), serving as an index of correct anticipation of the next character (Fig. 2a). We excluded trials involving rare transitions (20%) from the analysis because participants did not consistently exhibit optimal behaviour (i.e., ‘rare-change’) on these trials even if they had learned the current transition structure, because it also requires accurate knowledge about each character’s preference. In some cases, participants may have failed to change their initial prediction simply because they did not realise that the unexpected character had a different goal preference from the character they had anticipated, or because they were uncertain and chose not to change, given the potential cost associated with changing. In contrast, persisting during a paired transition suggests that participants correctly anticipated the next character and *vice versa*. Because whenever they correctly anticipated the paired character during the initial prediction, there were no other incentives for them to change. To ensure meaningful initial predictions rather than guesses, trials in which characters were previously observed no more than once and trials in which participants failed to make initial predictions within the time limit were excluded from the analysis (Fig. S3b). The “pair *history*_*t*_” regressor was defined as the number of occurrences of the paired transition in the last three trials from trial *t*. In the Supplementary Material, we indicate that using alternative reasonable history lengths yielded comparable results (Fig. S3d-f).

#### LM2

To test the influence of two types of pair history on participants’ decisions to persist, we conducted a second linear model (LM2), including both same-pair history (“experience”) and other-pair history (“inference”) as regressors:

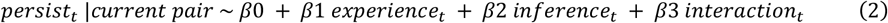

Specifically, if the current trial involved a *C*-to-*D* paired transition (Fig. 2a), the “experience” regressor was defined as the number of occurrences of the *CD* paired transitions (including *C*-to-*D* or *D*-to-*C*) in the last three trials. In contrast, the “inference” regressor corresponded to the number of occurrences of the *AB* paired transitions in the last three trials. Notably, “experience” and “inference” regressors were not anticorrelated, as there were also rare transitions (Fig. S2b). An interaction term between the experience and inference regressors was also included in the regression to ensure that behaviour was modelled robustly. However, this interaction was neither of primary interest nor statistically significant and thus is not discussed further.

#### LM3

In a third linear model (LM3), we modelled participants’ final goal preference predictions based on the goal history of the current character, similar to LM1:

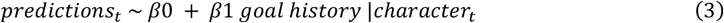

The participants’ final *predictions*_*t*_ about goal preference was modelled as left (0) or right (1), and the goal history counts the number of right shots from the current character in the last three times that this character was observed.

For all regression analyses, different periods (trial window lengths) were selected based on the specific analytical objective. Longer periods were used for analyses encompassing all trials, while shorter windows (typically the first 15 trials after a fresh start or reversals) were employed to isolate behaviour during the early learning period within each block. Different window lengths yielded consistent regression results in the fMRI experiment (Fig. S4b). Importantly, a minimum window length of 15 trials was required to ensure enough data points for reliable regression estimates.

#### Bayesian statistical analysis

One challenge of interpreting any null result in an fMRI experiment is that it might be due to a lack of statistical power in the fMRI data set (an absence of evidence) rather than the absence of an effect (evidence of absence). To distinguish between these possibilities, we employed Bayesian statistics. Bayes Factors are an established tool to assess whether the lack of a significant result is rather due to no effect (i.e. evidence of absence) or a lack of evidence (i.e. absence of evidence)^72^. We computed Bayes Factors (BF) for our key non-significant results using ‘Bayes Factor’ package in R. We used the categorisation to interpret our BFs.

### Computational modelling

We implemented three reinforcement learning (RL) models to capture how participants learned and used the task structure: a *Frequency Model* tracking character frequency, a *Transition Model* learning observable character-to-character (state-to-state) transitions, and a *Structure Model* incorporating both experienced and inferred transitions based on latent graph structure knowledge. All models shared the same goal-direction learning component and were fit to participants’ behavioural data using maximum likelihood estimation, with model comparison and simulation used to determine the best-fitting model.

#### Frequency Model

The *Frequency Model* only monitors how frequently each state (i.e., character) was observed and utilises this information to make predictions. In this model, four estimates [*F*(*A*), *F*(*B*), *F*(*C*), and *F*(*D*)] track the appearance frequency of each state. On each trial, the model updated the frequency of all states (ensuring that they always summed to 1) depending on whether they were observed or not (equation 4 for observed and equation 5 for other unobserved states). For example, if the observed state in trial *t* is *A*, the value updates will be computed as follows:

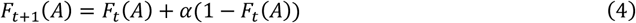

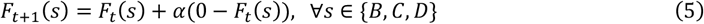

All frequency estimates were initialised at 0.25 and updated using a delta rule^25^, scaled by a learning rate *α* between 0 and 1.

#### Transition Model

The *Transition Model* estimates the transition probability between every two states (i.e., characters). It constructs a 4-by-4 transition matrix, wherein each element, denoted as *T*(*s* → *s*^′^), quantifies the probability of transitioning from a current state (*s*) to the subsequent state (*s*^′^). For example, *T*(*A* → *B*) indicates the transition probability from state *A* to state *B*. Therefore, the whole transition matrix looks like the one below:

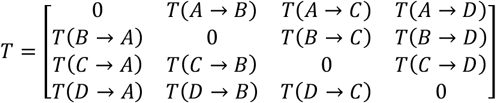

In this matrix, rows and columns represent the starting and ending states, respectively, and all self-to-self transitions are prohibited by setting *T*(*s* → *s*^′^) = 0 if *s* = *s*^′^. Besides, such a transition matrix adheres to the defining characteristics of a Markov chain, where the outgoing probabilities from each state sum to one: ∑_*s’*_ *T*(*s* → *s*^′^) = 1. Initially, all transition probabilities were set to 1/3, except for self-transitions, which were set to 0 and remained unchanged. Transition probabilities were updated using a simple update rule^25,27^, scaled by a transition learning rate *α* between 0 and 1:

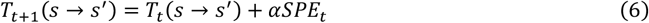

Where *SPE*_*t*_ indicates the state prediction error and is calculated for the transition currently experienced as follows:

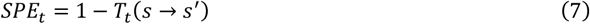

For example, if the learner observes a transition from *A* (trial *t*-1) to *B* (trial *t*), the equations will be as follows:

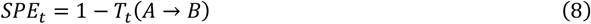

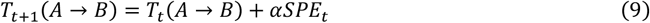

At the same time, the estimated transition probabilities for all states not reached (i.e., for all states *s*^′′^ other than *B*) are lowered accordingly, ensuring all the transition probabilities leaving from state *A* add up to one:

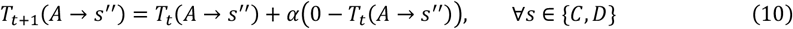

The probabilities of all other transitions not from *A* will remain unchanged.

#### Structure Model

The *Structural Model* extends the *Transition Model* by utilising structural knowledge to update both experienced transitions and inferred, unexperienced transitions. For example, if the current transition is from *A* to *B*, the model will not only update *T*(*A* → *B*) but also its bidirectional counterpart *T*(*B* → *A*). In addition, it updates the transition probability between *C* and *D* (i.e., *T*(*C* → *D*) and *T*(*D* → *C*)). By doing so, *T*(*A* → *B*), *T*(*B* → *A*), *T*(*C* → *D*), and *T*(*D* → *C*) are constantly maintained at equal values and are adjusted simultaneously. For simplicity, we denote these transition probabilities as a single variable *T*(*AB* & *CD*), reflecting the probability of the latent pairing structure being *AB & CD*. Instead of learning each transition independently, the *Structure Model* learns the latent pairing structure through observations, estimating probabilities for just three possible pairings: *T*(*AB* & *CD*), *T*(*AC* & *BD*), and *T*(*AD* & *BC*), simplified as *T*(*I*), *T*(*II*), and *T*(*III*), respectively, resulting in the following transition matrix:

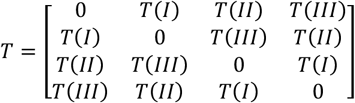

For example, if the learner observes a transition from *A* (trial *t*-1) to *B* (trial *t*), the model will compute a state prediction error (SPE) and update the probability of the corresponding pairing structure *T*(*AB* & *CD*), denoted by *T*(*I*). Critically, the model does the same update if a transition between *C* and *D* is observed. Updates are scaled by a transition learning rate *α* between 0 and 1, as follows:

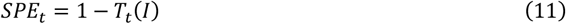

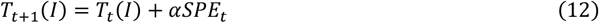

In the meantime, the probabilities of other pairing structures are updated too:

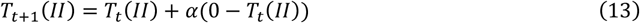

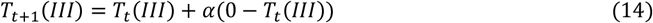

#### Goal-preference learning

The learning of each character’s goal preference is identical in all models: the models track four estimates [*G*(*A*), *G*(*B*), *G*(*C*), and *G*(*D*)], each indicating the expected goal preference of a given state (i.e., character). The update is performed using the same delta-rule model^25^. All *G*-estimates are initialised at 0.5 in the first trial. In every trial, the *G*-estimate of the observed character is updated based on its discrepancy from the actual direction outcome (0 or 1 for left or right, respectively), scaled by a goal-direction learning rate *η* between 0 and 1. The rest of the characters’ *G*-estimates remain unchanged. For instance, if character *A* appears in trial *t*, then its goal preference prediction error (GPE) and value update will be calculated as follows:

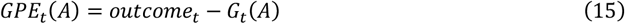

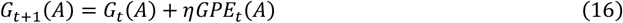

Assuming learners always act to maximise expected reward, the reward prediction error (RPE) equals the GPE when the action is rewarded (reward = 1), and equals the negative of GPE when the action is unrewarded (reward = –1), computed as:

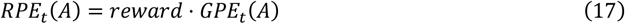

#### Action selections

Subsequently, each model integrates the information about goal preference and character transition to calculate the *Q*-values (see below) for the potential actions of making a save to the left or the right. Two actions will be taken in each trial: an initial prediction about the goal direction before knowing the character (*a*_*init*_) and a final prediction deciding whether to change it after seeing the character (*a*_*final*_).

#### Initial prediction

For the initial prediction, the agent has to assess which of the four states (characters) is the most likely to appear next and which direction this character prefers. The prediction is therefore the product of two factors: each character’s probability of appearing on this trial, and its potential goal direction. Here, the probability of a given state (*s*) appearing next is depicted by frequency estimates *F*(*s*) in *Frequency Model* or transition probabilities *T*(*s*) in the *Transition* and *Structure Models*. As before, the expected goal direction is represented by *G*(*s*) (1 indicates a rightward shot, and 0 indicates a leftward shot). For example, under the framework of *Structure Model*, the initial predicted possibility of the goal direction being right is ∑_*s*_ *T*(*s*)*G*(*s*), and the possibility of a leftward shot is ∑_*s*_ *T*(*s*)(1-*G*(*s*)). Considering the rule, where a correct prediction results in a 2-point gain while an incorrect one incurs a 2-point loss, the *Q*-values for predicting *Right* in any given trial *t* can be calculated as:

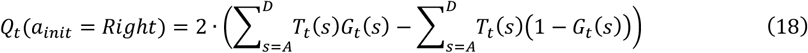

The *Q*-values for predicting *Left* is:

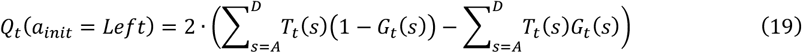

The *Q*-values of both actions are then used to determine the probability of predicting *Right* through a SoftMax function:

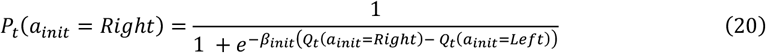

Here, the parameter *β*_*init*_ is the inverse temperature, which reflects the degree of stochasticity in the choice.

#### Final prediction

After seeing the character, an agent needs to make a final prediction, either changing the initial prediction or not. This choice is primarily based on the estimates of the current character’s goal preference *G*(*s*). For example, if it is character *A*, the potential expected goal direction will be *G*(*A*) (1 means rightward and 0 means leftward). The final prediction also depends on the initial prediction made, because there is a cost of changing or missing the initial prediction (0 points if correct, -2 points if incorrect) compared to persisting (+2 points if correct, -2 points if incorrect). Suppose the initial prediction (*a*_*init*_) is *Left*, and the current character is character *A*, the *Q*-value of changing it to *Right* as the final prediction is:

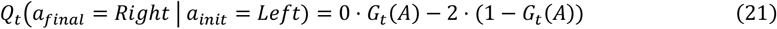

the *Q*-value of persisting with *Left* as the final prediction is:

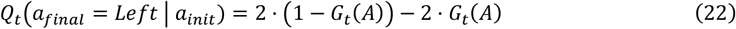

The probability of predicting *Right* as the final prediction (*a*_*final*_) is determined through a SoftMax function, with *β*_*final*_ as the inverse temperature parameter:

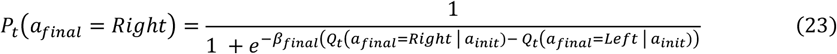

#### Model fitting

We used the maximum likelihood estimate (MLE) method to estimate each participant’s parameter. The log-likelihood (*LL*) was calculated as follows:

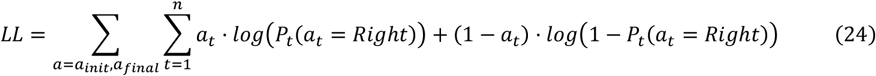

In this formula, *t* indicates the trial index, *n* represents the total number of trials. The actual action participants took is denoted by *a*, including the initial prediction *a*_*init*_, and the final prediction *a*_*final*_. If the participants chose *Right* in trial *t, a*_*t*_= 1, otherwise *a*_*t*_= 0. *P*_*t*_(*a*_*t*_*=Right*) is the probability of choosing the *Right* generated from a SoftMax function. The log-likelihood of each action is calculated as the sum of the log-likelihood values of all the unmissed trials. The final value of log-likelihood (*LL*) is the sum of log-likelihood of both actions (i.e., initial prediction and final prediction). We performed MLE using *optim()* function in R. We chose the ‘L-BFGS-B’ (limited-memory Broyden–Fletcher–Goldfarb–Shanno algorithm for bound-constrained optimisation) as the optimisation algorithm. We estimated four free parameters for every participant: state learning rate *α*, goal-direction learning rate *η*, and two inverse temperatures *β*_*init*_ and *β*_*final*_. For learning rates *α* and *η*, the upper and lower boundaries were 0 and 1, respectively, and the initial value was drawn from a uniform distribution between 0 and 1. For inverse temperatures *β*_*init*_ and *β*_*final*_, the lower boundary was 0, the upper boundary was infinite, and the initial value was drawn from a uniform distribution between 0 and 10. The maximum iteration limit is set to 10,000 times each run. We selected the optimal parameter set with the highest log-likelihood among 50 runs.

#### Model comparison

For model comparison, we used the Variational Bayesian Analysis (VBA) toolbox23 in MATLAB R2023b to compare the models using a random-effects approach. Given that all models contained the same number of free parameters, we used log-likelihood as the model evidence to compute the exceedance probability. This measures the likelihood of a particular model being the true model compared to the others.

#### Model simulation

We employed all three models to simulate synthetic data. For all models, inverse temperatures *β*_*init*_ and *β*_*final*_ were fixed at 1, the transition learning rate *α* (in equations 4, 5, 6, 9, 10, 12, 13, and 14) was set to 0.3 (with Gaussian noise = 0.05), and the goal-direction learning rate *η* (equation 16) was set to 0.25 (with Gaussian noise = 0.05). These parameter values were chosen based on the average of participants’ fitted parameters (Fig. S4f). We also performed parameter recovery and found that the Structure Model reliably recovered both learning rates most of the time (Fig. S4g). For each model simulation, it performed one of the tasks that the participants had previously encountered. We generated 120 simulated results for each model to ensure robustness. In these simulations, a model executed two actions per trial: an initial and a final prediction. The choice of every action was randomly drawn from a Bernoulli distribution with the probability of choosing right being *P*_*t*_(*a*_*t*_*=Right*) and the probability of choosing left as 1-*P*_*t*_(*a*_*t*_*=Right*). The simulated data were analysed using similar regression approaches to those used for human behavioural data.

### fMRI data analysis

#### Preprocessing

The fMRI data was preprocessed using FEAT, part of FSL version 6.00, from FMRIB’s Software Library^69,70^. For most of the analyses, data for each of the four blocks were pre-processed separately, except for the cross-block decoding analysis, where data from all blocks were concatenated prior to preprocessing. We used the default options in FSL: geometric distortion correction using B0 field mapping (effective echo spacing = 0.275 ms); motion correction using MCFLIRT; slice-timing correction for the interleaved sequence; brain extraction using BET^73^; spatial smoothing using a Gaussian filter kernel of FWHM 5.0 mm, and high-pass temporal filtering with 3 dB cut-off of 100 s. Registration of EPI images to high-resolution structural images and to standard (MNI) space was conducted using FMRIB’s Linear and Non-Linear Registration Tool, FLIRT and FNIRT^74^.

#### Whole-brain univariate analysis

We conducted three-level whole-brain statistical analyses using FSL’s FEAT^75^. The data were pre-whitened using FILM before the analyses to eliminate temporal autocorrelations^76^. At the first level, we applied a univariate general linear model (GLM) approach to each block separately for each participant. All whole-brain GLMs included four constant regressors: three of them represented the onset of every phase in a trial: the prediction phase, the character phase, and the outcome phase; another one represented the onset of motor response, including all the key presses. These constant regressors were set as 0-second spikes with amplitudes of one. Each GLM has one regressor of interest.

#### GLM 1

state transition regressor (onset: character phase, duration: 1.5 seconds, amplitude: ‘rare’ as 1 and ‘pair’ as -1; excluding the first four trials);

#### GLM 2

outcome/reward regressor (onset: outcome phase, duration: 2 seconds, amplitude: ‘correct’ as 1 and ‘wrong’ as -1, regardless of the actual points);

#### GLM 3

state prediction error (SPE) regressor (onset: character phase, duration: 1.5 seconds, amplitude: trial-by-trial estimates of SPE from Model 2; without the first four trials);

#### GLM 4

reward prediction error (RPE) regressor (onset: outcome phase, duration: 2 seconds, amplitude: trial-by-trial estimates of SPE from Model 2).

All regressors were convolved with the hemodynamic function. In all GLMs, the BOLD signal was denoised by including the six motion parameters derived from MCFLIRT realignment as the nuisance regressors. In the second level analysis, we averaged the estimates of the regressor of interest across four blocks using a fixed-effect approach for each participant. In the third level analysis, we estimated the group-level average using a mixed-effects approach (FLAME 1+2), accounting for the variance across participants^77^. The final results were obtained using whole-brain cluster correction with a voxel inclusion threshold of Z=4 and a cluster significance threshold of *P*<0.05.

#### ROI analysis

For each ROI analysis, we first defined a mask in the standard MNI space and then warped it to each subject’s individual structural space using FMRIB Software Library’s (FSL) non-linear transformations (FNIRT)^69^. Anatomical masks were used for ROIs whose locations were either supported by theoretical relevance or closely matched univariate activation patterns. These included: the hippocampus, defined using a probabilistic mask from a previous study^78^; the ventral striatum, from the Oxford-Imanova Striatal Structural Atlas^79^; the entorhinal cortex (EC), from the Juelich Histological Atlas^80^. Functional masks were created for regions with task-related univariate effects not well-aligned with anatomical boundaries. These were defined as 4-mm-radius spheres centred on peak activation coordinates with a slight adjustment to ensure coverage: OFC (MNI x=-28, y=36, z=-12), vmPFC (MNI x=-2, y=48, z=-10), and STG (left MNI x=-60, y=-16, z=6; right MNI x=64, y=-16, z=6). Due to their elongated anatomical shapes, masks for the insula and lateral prefrontal cortex (lPFC) were derived directly from whole-brain univariate maps, thresholded at Z = 4. All ROIs, except for vmPFC and OFC, were defined bilaterally and contained a sufficient number of voxels for multivariate analyses.

#### Multivariate analysis preparation

For multivariate analysis, we first extracted trial-specific beta estimates using a whole-brain GLM. We constructed a design matrix for each trial that was modelled with a separate regressor, time-locked to the onset of the event of interest (e.g., initial prediction, character onset, or outcome onset). We also included additional control regressors: two regressors for the remaining non-interest phases (collapsed across trials), one regressor for all motor responses, and six motion parameters (generated from motion correction, as described in “Preprocessing”) as nuisance regressors. For instance, when analysing character onset, the model included *n* trial-specific regressors time-locked (one per trial, *n* equals the number of trials), a single regressor each for the initial prediction and outcome phases, and other control regressors. GLMs were done by Matlab (2023b) and Statistical Parametric Mapping software (SPM12; Wellcome Trust Centre for Neuroimaging, London, UK).

#### Representational similarity analysis (RSA)

We used representational similarity analysis (RSA)^26,81^ at the trial level to test whether the ROIs carried information about task variables. Trial-level RSA was performed within each block by employing the following steps:

1. Constructing the model representational dissimilarity matrix (RDM). The *relational (transition) structure* RDM inversely mirrors the transition probability matrix, wherein each cell value is calculated as one minus the transition probability. For example, in a block with AB & CD pairs, if the character in trial *t* is A and trial *t*+1 is B, the dissimilarity value should be one minus the transition probability between A and B (paired transition): 1 - 0.8 = 0.2. If the character in trial *t* is A and trial *t*+2 is D, the dissimilarity value should be one minus the transition probability between A and D (rare transition): 1 - 0.2 = 0.8. A dissimilarity value of 1 (highest) is assigned when characters do not occur consecutively across trials (never transition). Notably, all the pairings with the same character (‘self’) were excluded from the analyses. Similarly, the *goal preference prediction RDM* was constructed by comparing the predicted direction (e.g., left or right) across all trials. Each cell was assigned a value of 0 if the predicted goals for both trials were the same (e.g., left– left or right–right), and 1 if they differed (e.g., left–right). Additionally, the first four trials were omitted from the RDM calculations because participants needed a few trials at the start to learn.
2. Calculating neural RDM within ROIs. This RDM was constructed using the Mahalanobis distance (Euclidean distance after multivariate noise normalisation) between the activity patterns for all pairings allowed by the model RDM^82^. Depending on the analysis, we used beta estimates time-locked to character onset, outcome onset, or initial prediction onset.
3. Computing the correlation between model RDM and neural RDMs. We first calculated the rank correlation (Kendall’s tau) between model RDM and neural RDMs of all ROIs for each block and then averaged the correlations across blocks for each participant. We used the Wilcoxon signed-rank for significance. The resulting p-values were corrected for multiple comparisons across ROIs using the Holm–Bonferroni method^83^.

The same steps were used for all the analyses: *relational (transition) structure* (time-locked to character and outcome onset), and *relational (transition) structure* of anticipated characters based on paired transitions from the previous character (time-locked to initial prediction onset). In a further set of follow-up RSA analyses time-locked to outcome onset, data were separated into two parts based on the type of reward received (correct vs. wrong), and each part of the data was analysed independently.

Notably, this trial-level RSA approach controls for confounds from fMRI autocorrelation, as paired characters often occur in adjacent trials, by ensuring that comparisons between trials of the same transition type yield consistent values, regardless of their temporal proximity. For example, in a block where *A* and *B* are paired, if the character of trial *t*=5 is *A*, trial *t*=6 is *B* (adjacent), and trial *t*=36 also features *B* (distant), the dissimilarity between trials *t*=5 and *t*=6 is the same as between *t*=5 and *t*=36 (e.g., 0.2), despite their differing temporal proximity.

#### RSA with partial correlation

We performed partial correlation analyses to control for potential confounding variables. For example, when testing the correlation between the *goal preference prediction* RDM and neural RDMs, we controlled for the influence of the *dense relational structure* RDM by regressing out its variance from both prior to the correlation calculation. The *dense relational structure* RDM is a *relational structure* RDM that does not exclude self-comparisons, because *goal preference prediction* RDM allows comparisons between trials with the same character. Unlike the sparse *relational structure* RDM mentioned above, the *dense relational structure* RDM retains all cells, including those with a value of 0 (“self”). This approach was applied to all analyses requiring control for competing models: (1) testing hypothetical *relational structure* (initial prediction epoch) while controlling for *initial goal preference prediction*, (2) testing *relational structure* (character epoch) while controlling for *goal preference prediction*, and (3) testing *relational structure* (outcome epoch) while controlling for *actual goal direction*.

#### Whole-brain searchlight RSA

We conducted a whole-brain searchlight RSA to identify the task’s relational (transition) structure representation in the brain. We defined a sphere (radius = 20 mm) containing approximately 100 voxels around each searchlight centre voxel. The neural RDMs were calculated for each sphere, and behavioural model RDMs for relational (transition) structure were created using the procedure explained above (see “Multivariate analysis preparation” and “Representational similarity analysis (RSA)” for details). Again, we used rank correlation (Kendall’s tau) to quantify the extent to which pattern activity within each searchlight was explained by the model RDM. This yielded a Kendall’s tau correlation coefficient for each searchlight position, reflecting how well local neural activity patterns encoded the relational structure. We then transformed the correlation maps (Kendall’s tau) to z-scores for statistical normality using a Fisher z-transform. The single-subject Z-transformed correlation maps were registered to a MNI standard space using FSL’s applywarp. For group-level significance test, we used FSL’s nonparametric permutation test tool, randomise (5000 permutations), with the Threshold-Free Cluster Enhancement (TFCE) option to correct for multiple comparisons^84^.

#### Principal component analysis (PCA)

We performed principal component analysis (PCA) on trial-specific fMRI beta estimates to visualise the neural geometry of the *relational (transition) structure* representation. Beta estimates were obtained for each trial within each participant’s block and ROI (see “Preparation for the multivariate analysis”). Before PCA, the data were normalised using Min-Max scaling (rescaling voxel activations between 0 and 1) separately for each trial. The PCA was conducted at the block level using the PCA package (https://pypi.org/project/pca/). The voxel-wise beta patterns were projected onto a three-dimensional PCA space, defined by the first three principal components explaining the most variance. We then computed the mean PCA coordinates for each character by averaging across trials involving the same character, controlling for temporal proximity. Connections between characters were visualised according to the pairing structure of that block: paired characters were connected with thicker lines, while unpaired ones were connected with thinner lines, providing an intuitive neural representation of the relational structure.

#### Decoding analysis

We performed multivariate decoding analyses using support vector machine (SVM) classification with a linear kernel in Python using the scikit-learn library (https://scikit-learn.org/stable/)^85^. For each participant, trial-level beta estimates were extracted from predefined regions of interest (ROIs) using a GLM time-locked to specific task events (e.g., character onset). Before decoding, beta estimates were normalised using Min-Max scaling separately for each trial. Decoding was performed using repeated cross-validation (20 iterations), randomly splitting data into training (75%) and testing (25%) sets in each iteration. Classification accuracy was averaged across iterations for robustness. The decoding pipeline was implemented separately for each block, participant, and ROI, with labels derived from experimental design or data (e.g., goal prediction). For goal direction decoding, predicting a leftward goal direction was labelled as 0, whereas predicting a rightward direction was labelled as 1. To mitigate potential confounds arising from temporal autocorrelation in fMRI data, where consecutive trials may exhibit correlated noise due to the temporal proximity of paired characters, we included only blocks where paired characters’ preferred goal directions are the opposite (i.e., labelled differently). This approach reduces the risk of inflated similarity in neural representations that could otherwise bias decoding analyses. Each participant’s decoding accuracy was then calculated by averaging results across all relevant blocks. Because decoding accuracies were normally distributed across participants in all ROIs (Shapiro–Wilk test, all W>0.95, all *P*>0.05), we used paired *t*-tests for significance testing. The resulting p-values were corrected for multiple comparisons using the Holm– Bonferroni method^83^.

#### Cross-condition generalisation decoding analysis

To evaluate the generalisation of neural representations across task conditions, we conducted cross-block decoding analyses. For cross-pair goal prediction decoding, the neural data and labels were selected and normalised as described above, then grouped according to character pairs. For example, in a block with AB & CD pairs, neural data from trials involving characters A and B were combined into one set, while trials with characters C and D formed the other set. Given our selection criteria (blocks in which paired characters had opposite preferred goal direction), both sets contained balanced goal predictions (left and right). Classifiers were trained to decode goal preference predictions from neural data of one character pair (e.g., AB pair) and tested on the other pair (e.g., CD pair), and vice versa. Cross-pair decoding accuracy was averaged across both training-testing directions to ensure unbiased estimates. Decoding accuracies were computed separately for each block and then averaged across blocks for each participant. For cross-block context decoding, classifiers were trained to decode contexts on neural data from one set of blocks (e.g., blocks 1 and 2) and tested on another set (e.g., blocks 3 and 4), and vice versa. To enable accurate cross-block comparison, neural data from all blocks were concatenated prior to preprocessing. For each participant and ROI, normalization (Min-Max scaling) was applied independently for training and testing sets. Decoding accuracy for each participant and ROI was calculated separately for each cross-block pairing and averaged across both training-testing directions.

All decoding results were assessed at the group level for each ROI. Significance testing involved comparing decoding accuracies against empirical chance levels (e.g., 50% for goal preference predictions) using one-tail t-tests.

#### Time-course analysis

Time-course analyses were performed using MATLAB. We separated the time-course data into epochs through the following steps: (1) Extracting the filtered BOLD time-course data from each voxel within the ROI (individual space) and averaging across voxels; (2) Normalising the average time-course data; (3) Up-sampling the normalised data by a factor of 10; (4) Interpolating the up-sampled data using the polynomial form of the cubic spline method in MATLAB; (5) Epoching the interpolated data into multiple 10-s windows, each aligned to the onset of the regressors of interest. Regressions were then fitted to the epoched data at each time step using the ordinary least squares method. For all time course analyses, the parameter estimates at each time step were averaged across all trials for every participant using a fixed-effects approach. Statistical significance at the group level was evaluated at each original time point corresponding to each TR of the original fMRI data, rather than the interpolated data. Given the results from different time points were not independent from each other, the resulting p-values were corrected for multiple comparisons across time points using the false discovery rate (FDR) with Benjamini-Hochberg (BH) procedure^86^. The peak p-values were corrected for multiple comparisons across ROIs using the Holm–Bonferroni method^83^.

### Neural network model

#### Neural Network Architecture

We employed a recurrent actor-critic network implemented in PyTorch^87^. The agent’s policy and value function share a single-layer LSTM core (64 hidden units) followed by separate output heads. At each time step, the LSTM receives a 10-dimensional input vector including trial number *t*, the current state (*s*_*t*_), the previous action (*a*_*t*−1_ | *s = s*_*t*_) and reward (*r*_*t*−1_ | *s = s*_*t*_) when the same state was encountered before. The current state and the previous action were coded as 4-dimensional one-hot vectors. a 64-dimensional hidden state. The actor’s head is a fully connected layer mapping the LSTM hidden state to 4 action logits (one per possible action/state), followed by a SoftMax to output a probability distribution over actions. The critic’s head is another linear layer that outputs a scalar value estimate for the current state. Specifically, the LSTM cell dynamics follow the standard equations^88^:

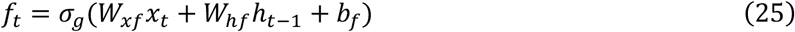

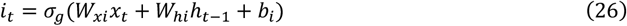

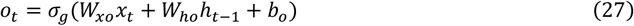

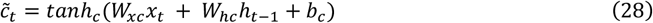

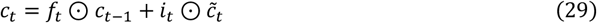

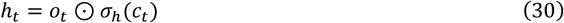

Here, *x*_*t*_ denotes the input vector at time *t*; *h*_*t*−1_ is the previous hidden state, and *σ* is the logistic sigmoid activation function. The gates *f*_*t*_, *i*_*t*_, *o*_*t*_, represent the forget/maintenance gate, input gate, and the output gate, respectively. 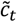 is the candidate memory content, and ⊙ denotes element-wise multiplication. These dynamics allow the network to maintain and selectively update memory traces over time, enabling the agent to accumulate information across a long time scale.

#### Training algorithm

Training employed an Advantage Actor-Critic (A2C) algorithm within a meta-reinforcement learning (meta-RL) framework based on the RL^2^ paradigm^89^. The agent learned to quickly adapt to a distribution of tasks drawn independently and identically (i.i.d.), without explicit task identifiers. Fast within-episode adaptation occurred solely through changes of the hidden units’ activities, while slower across-episode learning was achieved via gradient-based updates to the network weights. After each episode, the policy *π* (parameterized by neural network parameters *θ*) was updated using a policy-gradient loss, computed as the negative log-probability of the selected actions weighted by the advantage *A*_*t*_, and a value loss, calculated as the mean-squared error between the predicted value and the actual return. The total loss function was:

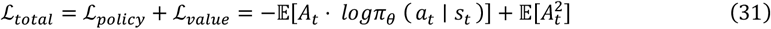

Advantages *A*_*t*_ were computed as:

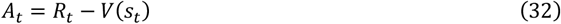

with discounted, bootstrapped returns defined as:

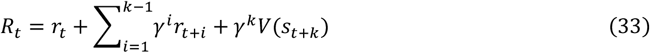

Here, *a*_*t*_, *s*_*t*_, and *R*_*t*_ represent the action, state, and discounted n-step bootstrapped return at time, respectively, *γ* is the discounting factor, and *k* is the number of steps until the episode ends and is upper bounded by the maximum unroll length.

Network weights, including LSTM kernels and output heads, were initialised using PyTorch defaults and trained end-to-end using gradient descent via the Adam optimiser. The learning rate was systematically tuned within the range [0.001, 0.01], typically set at 0.005. The discounting factor *γ* was also tuned within the range [0, 0.99], and typically set at 0, given no value/reward dependency across states. Training was strictly on-policy with an effective batch size of one episode per update, without employing replay buffers, dropout, batch normalisation, layer normalisation, or weight decay. Advantages were standardised per episode to stabilise training. Each model was trained for 5000 episodes, each comprising 160 trials. The LSTM hidden and cell states were reset to zero at the start of each episode to ensure within-episode adaptation. Training performance was monitored every 100 episodes by evaluating the model on 10 novel tasks and recording mean scores, accuracy, and losses to validate learning progress. Training was terminated after 5000 episodes when performance plateaued, without early stopping.

#### Environment details

The agent performed a character (state) prediction task in a four-state Markov decision process environment. At each trial, the agent received input of the current state, along with the previous action and reward encountered in the same state, and the current trial index (as mentioned above). The agent then predicted the next state, receiving a reward from the environment: +1 for a correct prediction and −1 otherwise. Each episode instantiated a distinct 4-state environment governed by a hidden transition matrix following the two-pair transition structure, with two paired transitions (0.8) and two rare transitions (0.2), with no self-transitions allowed. Each episode (160 trials) included two transition reversals—one occured randomly around midway through the first half and another in the second half—swapping paired and rare transitions. This required the agent to adapt to changing dynamics. The agent’s actions did not affect the actual state transitions; they solely determined the reward based on prediction accuracy. Unlike the *Goalkeeper Game*, this task removed directional prediction and focused entirely on learning latent relational (transition) structure from feedback. Under the fixed structural constraints, six distinct transition matrices were possible by permuting state pairings. A new matrix was randomly sampled in each episode to encourage generalisation and rapid adaptation to novel dynamics.

#### Behavioural assessment

Agent behaviours were assessed based on both prediction accuracy per episode and the learning strategies. Similar to the “Behavioural data analysis” that was performed on the human participants, we also analysed model behaviour by performing logistic regression to predict trial-by-trial accuracy based on recent transition history. Two primary regressors were defined: same-pair history (“experience”) and other-pair history (“inference”), using a history length of three trials. The dependent variable was the agent’s state-prediction accuracy in the current paired transition. Again, an interaction term was included for robustness but not discussed further as it was neither significant nor of primary interest. Different trial window lengths were selected based on the specific analytical objective. Longer windows were used for analyses encompassing all trials, while shorter windows (typically the first 15 trials) were employed to isolate behaviour during the initial learning period within each block.

#### Single-unit relational (transition) structure RSA analysis

We performed single-unit representational similarity analysis (RSA) on the trained neural network, analogous to the RSA applied to fMRI data. A theoretical representational dissimilarity matrix (RDM) was constructed at the trial level to capture the relational (transition) structure, where dissimilarity values were inversely related to the transition probabilities (see “Representational similarity analysis (RSA)” for details). For each hidden LSTM (short-term memory component) unit, we computed a neural RDM from its trial-by-trial activation patterns. Because each unit produces a single activation value per trial, the dissimilarity between two trials was defined as the absolute difference in that unit’s activation. The RSA score was calculated using rank correlation (Kendall’s τ) between neural RDM and model RDM. As each episode contained two latent relational (transition) structures due to reversal events, the RSA scores were calculated separately for each structure and averaged to obtain a per-episode score. This process was repeated across 40 episodes, and the resulting scores were averaged to yield a robust RSA value for each of the 64 hidden units. As a baseline, we repeated the same single-unit RSA procedure on an untrained (randomly initialised) network to estimate chance-level correlations as a null distribution.

#### Network lesion analysis

To assess the causal contribution of task-relevant units, we performed an artificial lesion analysis on the trained neural network agent. Units whose RSA scores significantly fall out of the null distribution (obtained from an untrained network) were classified as relational-structure-encoding, or “HPC-like,” units. In most trained networks, approximately half of the 64 hidden units met this criterion. The remaining units, whose RSA scores did not differ significantly from chance, were classified as “control units”. This yielded two equally sized groups of 32 units each. Lesions were implemented by setting the hidden-state outputs of selected units to zero during task performance, effectively silencing their contribution without altering the network’s learned weights. To quantify the functional impact of these lesions, we assessed changes in the agent’s learning strategy using the regression framework described in “Behavioural assessment,” focusing on experience- and inference-driven behaviour. We systematically varied the lesion percentage range from 0% to 100% of each group (in 10% increments). For each level, a random subset of the specified proportion (e.g., 50%) of units were randomly selected and silenced. The agent’s behaviour was then evaluated on a single episode. For example, when the lesion percentage was set to 50%, we randomly lesioned 50% of the HPC-like units and assessed the agent’s learning strategy in one episode, followed by randomly lesioning 50% of the control units in a separate episode. This procedure was repeated 10 times per lesion level, each time with a new random subset of units. Final performance scores were averaged across the 10 episodes to reduce noise. For each lesion level, this process was repeated for 120 runs, yielding 120 simulated data points per condition. Crucially, the network’s weights remained fixed during all post-lesion evaluations, allowing us to isolate the impact of silenced units and directly compare the behavioural consequences of lesioning HPC-like versus control units.

