## Supplementary material for "Causal necessity of human hippocampus for structure-based inference in learning": Fig. S

### Supplementary Figures

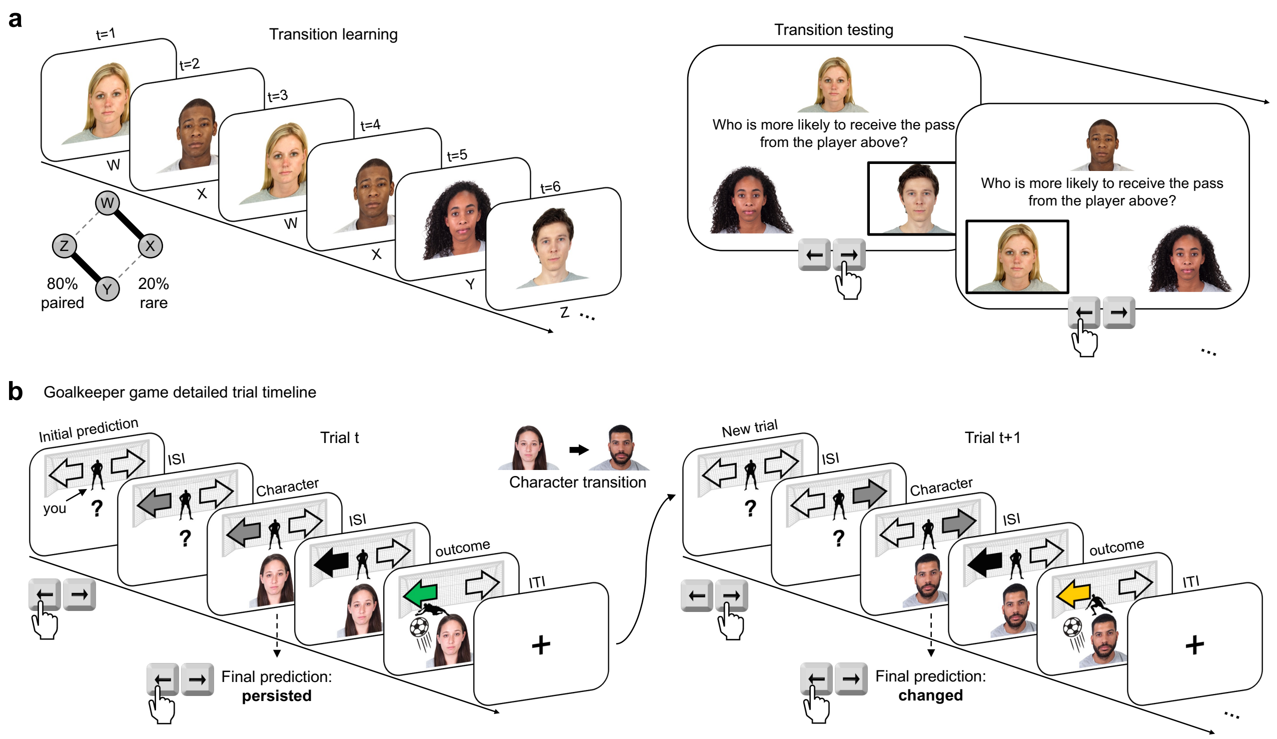


**Figure S1. Detailed trial timeline for the Transition Training and the Goalkeeper Game. a.** An example of a *Transition Training* block, including both the learning and testing parts. See "Transition training" in the "Experimental tasks" section of the Methods for details. This example shows a sequence with *WX* & *YZ* as two pairs (80% paired transitions), corresponding to the cartoon schematic shown in Fig. 1d. **b.** Two example trials from the Goalkeeper Game, shown as a detailed version of Fig. 1a. See "Goalkeeper Game (main task)" in the "Experimental tasks" section of the Methods for details. In this example, the participant initially predicted "Left", persisted with it, and received +2 points for a correct prediction on trial *t*; on trial *t*+1, they initially predicted "Right" but changed to "Left" in the final prediction, earning 0 points for a correct but changed response.

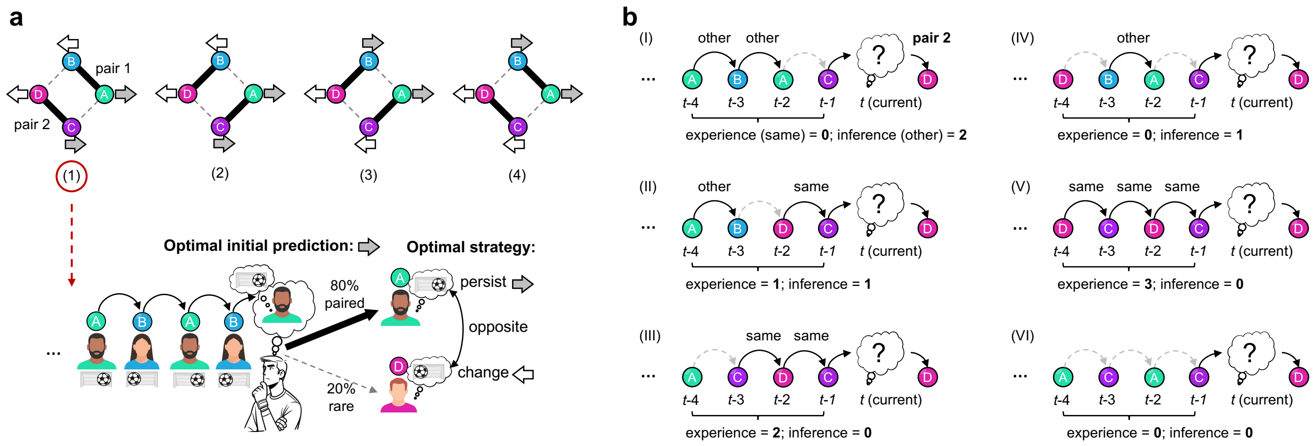


**Figure S2. Optimal behaviour and paired transition history. a.** The top panel shows four possible combinations of transition structure and characters' preferred goal directions. In each block, four characters form two pairs, and there are always two characters that prefer left and two that prefer right. Paired characters either have the same goal preferences (implying opposite preferences for rare transitions) or have opposite preferences (implying same preferences for rare transitions). Thus, any two characters linked to the same predecessor via different transition types always have opposite goal preferences. The bottom panel illustrates the optimal behaviour. During the initial prediction phase, participants should anticipate the next character based on paired transitions and persist with their initial predictions if that paired character indeed appears. For example, in a block with AB & CD pairs, where A and C prefer right and B and D prefer left, seeing B in trial t-1 should lead to an initial prediction of right in trial t by anticipating A (paired). Thus, they should persist if A indeed follows, or change to left if D appears instead (rare transition). **b.** Illustration of paired transition history. Given a certain current paired transition (e.g., C-to-D, pair 2), six possible scenarios can arise, differing in the number of recent same-pair transitions ("experience") and other-pair transitions ("inference"). Scenario (I) corresponds to the example shown in Fig. 2a.

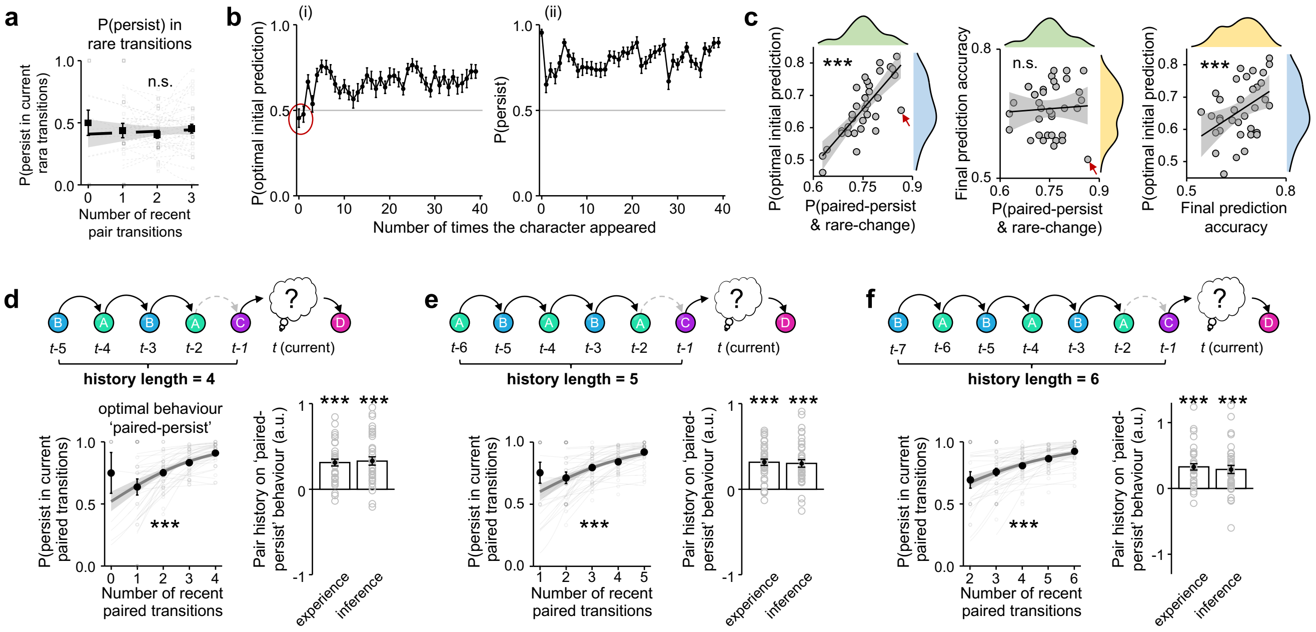


**Figure S3. Supplementary behavioural analyses of the fMRI experiment. a.** Results of linear mixed-effects models using rara transitions. The pair history does not significantly influence participants' likelihood of persisting in the current rare transition ($\beta$±CI=-0.082±0.356, *P*=0.817). Using another new linear mixed model (LM4) that involves both paired and rare transitions, $persist_t \sim pair\ history_t * current\ transition\ type\ (paired\ vs\ rare)$, we show that the influence of pair history on participants' likelihood of persisting is significantly higher when the current transition is a paired transition than a rare transition (interaction: $\beta$±CI=1.157±0.240, *P*=1.46×10$^{-6}$). This is because 'paired-persist' behaviour relies solely on understanding of the transition structure, whereas 'rare-change' behaviour also requires knowledge of character preferences, the influence on; therefore, only 'paired-persist' is significantly influenced by the pair history (main effect of pair history: $\beta$±CI=1.038±0.211, *P*=8.77×10$^{-7}$), but not the 'rare-change', which is excluded from our analyses. **b.** (i) The probability of making an optimal initial prediction is plotted against the number of times the current character appeared. An optimal initial prediction is defined as participants' correctly anticipating the next character based on paired transitions and making an initial prediction based on the anticipated character's preferred goal direction (see Fig. S2a). Performance was near chance when the character was observed no more than once (red circle), and such trials were excluded from the regression analyses. (ii) The probability of persisting with the initial prediction is plotted against the number of times the current character appeared. Participants tended to only persist at the beginning. **c.** Correlations among optimal initial prediction, optimal persist behaviour (paired-persist & rare-change), and final prediction accuracy. The optimal initial prediction and the optimal (transition-based) 'paired-persist' behaviour are positively correlated with each other (left, Pearson's *r*=0.784, *P*=1.54×10$^{-18}$). Optimal initial prediction required knowledge of both transition structure and each character's preferred goal direction, whereas optimal persist behaviour relied only on the transition structure, providing a cleaner measure of this knowledge. As expected, final prediction accuracy, reflecting goal preference knowledge, was not correlated with the optimal persist behaviour (middle, Pearson's *r*=0.784, *P*=0.718), but was positively correlated with the optimal initial prediction (right, Pearson's *r*=0.470, *P*=0.004). An example participant (indicated by the red arrow) demonstrated strong transition knowledge (optimal persist behaviour) but weak goal direction knowledge, resulting in a relatively low optimal initial prediction. Data distributions are shown at plot edges (blue: optimal initial prediction; green: optimal persist behaviour; yellow: final prediction accuracy). **d–f.** Results of linear mixed-effects models with different history lengths (4–6 trials). The top panels illustrate examples of different history window lengths. The left panels show that the more paired transitions participants encountered, the more likely participants were to persist in the current paired transition (History length=4: $\beta$±CI=0.317±0.040, *P*=2.08×10$^{-9}$; History length=5: $\beta$±CI=0.328±0.039, *P*=4.47×10$^{-10}$; History length=6: $\beta$±CI=0.332±0.038, *P*<10$^{-16}$). Each grey dot represents the data of an individual participant for a given pair transition history; thin light-grey lines show individual regression fits. Black dots with error bars show group mean ± SEM. The group-level regression fit is plotted as a thick, dark grey line with 95% CI (without random effect, for visualisation purposes only). The right panels show that both recent experience of same-pair history and inference from other-pair history significantly increased participants' likelihood of 'paired-persist' behaviour (History length=4: experience $\beta$±CI=3.045±0.370, $P<10^{-16}$; inference $\beta$±CI=3.042±0.447, $P=9.70\times10^{-12}$. History length=5: experience $\beta$±CI=3.167±0.336, $P<10^{-16}$; inference $\beta$±CI=2.945±0.424, $P=3.78\times10^{-12}$. History length=6: experience $\beta$±CI=3.611±0.478, $P=4.44\times10^{-14}$; inference $\beta$±CI=3.275±0.560, $P=5.06\times10^{-9}$). The history length must exceed two to ensure independence between experience and inference histories; however, a longer length means excluding more trials at the beginning. We therefore used a history length of three in the main analysis, as it yielded results consistent with those from longer history lengths. Grey dots represent the individual regression estimates. The bar plots indicate group beta mean ± SEM. ***$P<0.001$. n.s., not significant.

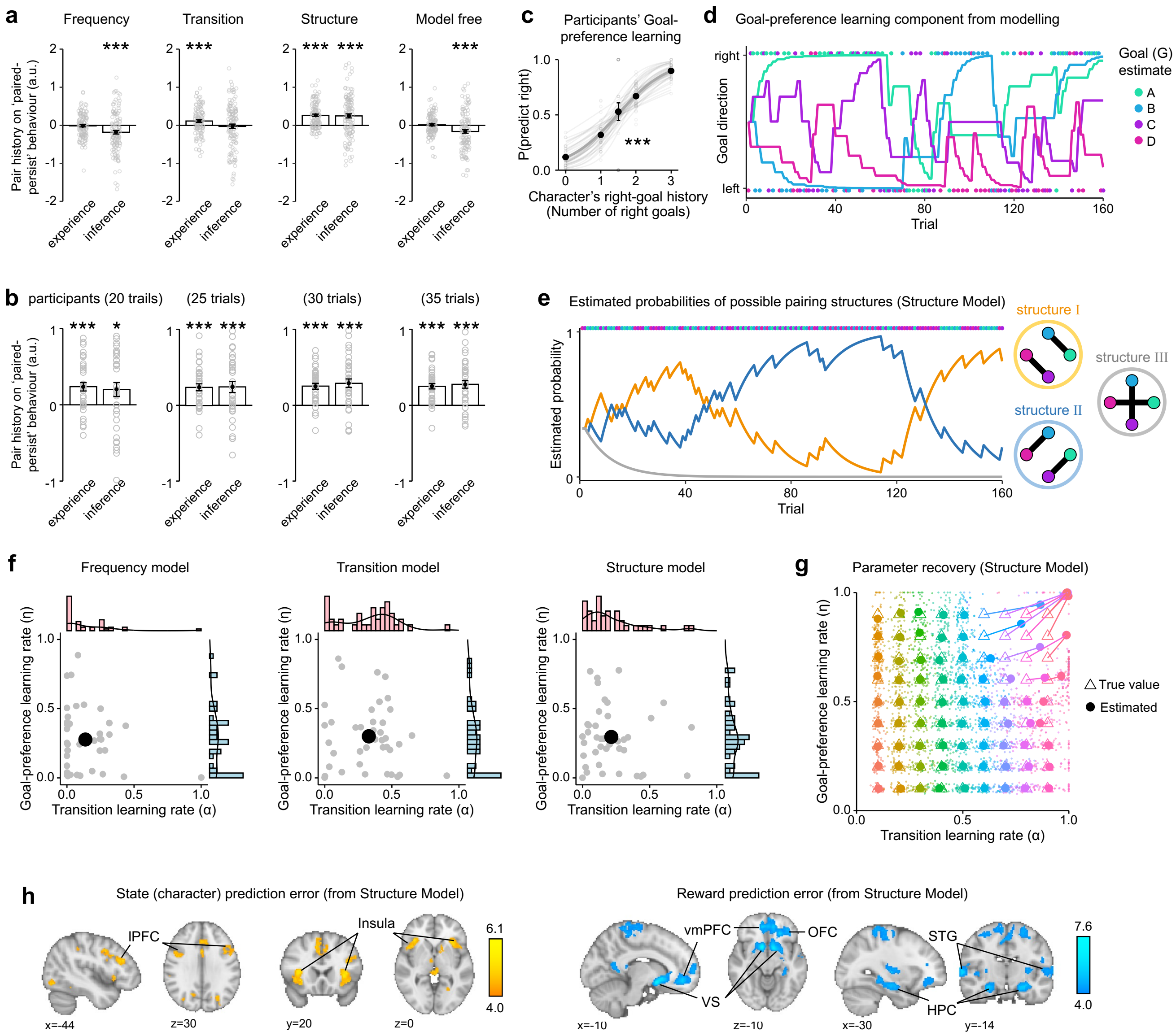


**Figure S4. Supplementary analyses of the behavioural and modelling results. a.** Regression analysis of paired-persist behaviour as a function of same-pair (experience) and other-pair (inference) history in model-simulated data during the learning period (first 15 trials per block). Consistent with data using all trials (Fig. 2g), during the learning period: the Frequency Model failed to learn from same-pair experience ($\beta$±CI=-0.024±0.145, $P$=0.866) and was misled by the other-pair transitions ($\beta$±CI=-0.771±0.249, $P$=0.002); the Transition Model relied only on the same-pair experience ($\beta$±CI=0.658±0.147, $P=7.29\times10^{-6}$), showing no inference from other-pair history ($\beta$±CI=-0.137±0.263, $P$=0.604). The Structure Model utilised both same-pair experiences ($\beta$±CI= 1.460±0.175, $P<10^{-16}$) and other-pair inferences ($\beta$±CI=1.245±0.266, $P=2.87\times10^{-6}$). We also examined an additional model-free approach and found that it replicated the Frequency Model's simulated behaviour. This model does not learn about character identities; instead, it makes initial predictions based solely on the overall frequency of goal directions (left vs. right), without anticipating which character will appear. Upon seeing the character, it decides whether to persist or change using the same goal-direction learning component as other models. As a result, this model-free approach generated a similar effect as the Frequency Model (all trials: experience $\beta$±CI=0.083±0.098, $P$=0.400; inference $\beta$±CI=-0.885±0.152, $P=6.40\times10^{-9}$; learning period (15 trials): experience $\beta$±CI=0.152±0.144, $P$=0.292; inference $\beta$±CI=-0.664±0.243, $P$=0.063). **b.** Robust effects of experience and inference across different trial windows in participant data. Regression analyses across various trial windows (periods = 20-35 trials) in the fMRI experiment showed that both experience of same-pair history and inference from other-pair history significantly increased participants' likelihood of 'paired-persist' behaviour (period = 20 trials: experience $\beta$±CI=1.608±0.345,

$P$=3.11×$10^{-6}$; inference $\beta$±CI=1.418±0.558, $P$=0.011; period = 25 trials: experience $\beta$±CI=1.718±0.280, $P$=8.68×$10^{-10}$; inference $\beta$±CI=1.760±0.430, $P$=4.26×$10^{-5}$; period = 30 trials: experience $\beta$±CI=1.938±0.278, $P$=3.00×$10^{-10}$; inference $\beta$±CI=2.113±0.420, $P$=4.86×$10^{-7}$; period = 35 trials: experience $\beta$±CI=2.070±0.277, $P$=7.45×$10^{-14}$; inference $\beta$±CI=2.137±0.390, $P$=4.27×$10^{-8}$). **c.** Participants learn to predict goal direction from the history: the probability of making a rightward final prediction increased with the number of times a character had previously shot to the right ($\beta$±CI=0.802±0.028, $P<10^{-16}$). Each grey dot represents the data of an individual participant for a given goal history; thin light-grey lines show individual regression fits. Black dots with error bars show group mean ± SEM. The group-level regression fit is plotted as a thick, dark grey line with 95% CI (without random effect, for visualisation purposes only). **d.** An example of the goal-direction learning component from one participant (see "Computational modelling: Goal-direction learning" in the Methods). The goal learning rate was estimated with the sequence learning component using the Structure Model. Colours indicate character identity, dots represent their actual goal directions (left=0 and right=1), lines show estimated goal-direction (G-estimate) for corresponding characters. **e.** Example of estimated probabilities for the three possible latent pairing structures from the same participant, as fit by the Structure Model (see "Structure Model" in Methods). The model tracks the probability of AB & CD (orange), AC & BD (blue), and AD & BC (grey), denoted as structures I, II, and III, respectively. **f.** The model-estimated state-transition learning rates (α) and goal-direction learning rates (η) across participants are plotted for three models separately. Distributions are shown at plot edges. The large black dots indicate the group average. **g.** Parameter recovery for the Structure Model. We assessed parameter recovery by simulating behavioural data using the Structure Model with systematically varied state-transition and goal-direction learning rates (0.1 to 0.9, all combinations). For each parameter pair, we generated 40 simulated datasets using sequences from the human experiment and fit the model to recover the underlying true values. True parameter values are shown as triangles, estimated values as dots; small dots represent individual simulations, and larger dots indicate the mean estimate for each parameter pair. The Structure Model accurately recovered learning rates except when both rates were very high, indicating strong overall reliability of the model's parameter estimates. **h.** Brain regions encoding state/character prediction error (Methods, GLM3; yellow) and reward prediction error (GLM4; blue) estimated from the Structure Model closely correspond to brain regions representing character transition and outcome processing (Fig. 3a), respectively. The colour bar indicates the Z-score. All plots are cluster-corrected using a threshold of Z > 4.

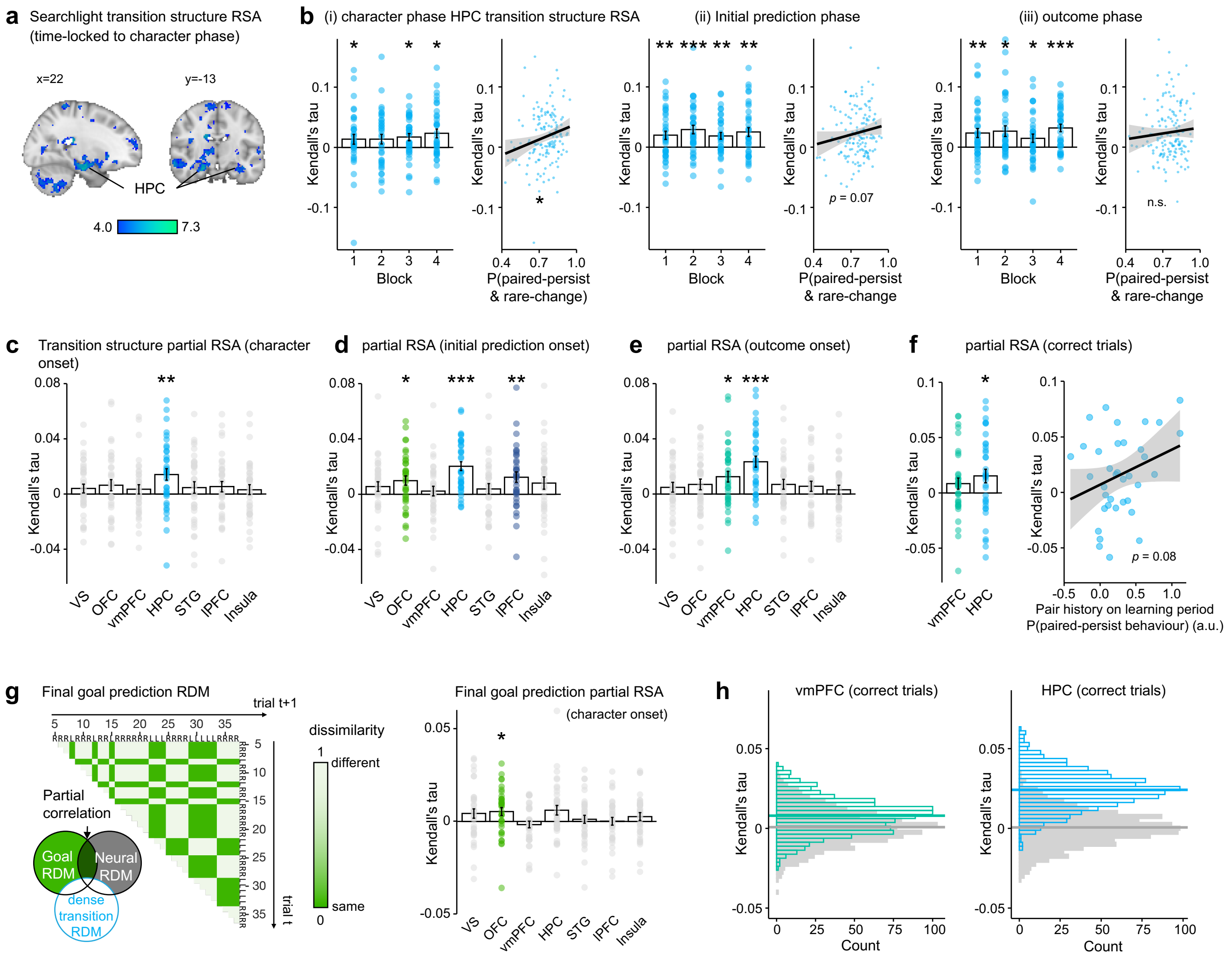


**Figure S5. Supplementary RSA analyses. a.** Whole-brain searchlight RSA identified representations of transition structure between characters (time-locked to character onset) in the hippocampus (HPC: left peak Z=6.21, MNI: x=22, y=-16, z=-22; right peak Z=5.06, MNI: x=-30, y=-16, z=-24). The colour bar indicates the Z-score, thresholded at Z > 4. **b.** Block-level transition structure RSA effects (rank correlations between model and neural RDMs; Kendall's tau) in HPC, shown separately for different event onsets: (i) character, (ii) initial prediction, and (iii) outcome. HPC consistently encoded transition structure across blocks. Given non-independence among blocks, p-values were adjusted using false discovery rate (FDR) correction: Significant HPC representations occurred at character onset (block 1, *V*=480, $p_{FDR}$=0.013; block 2, *V*=434, $p_{FDR}$=0.058; block 3, *V*=508, $p_{FDR}$=0.010; block 4, *V*=494, $p_{FDR}$=0.010), initial prediction onset (block 1, *V*=492, $p_{FDR}$=0.006; block 2, *V*=563, $p_{FDR}=6.23\times10^{-3}$; block 3, *V*=527, $p_{FDR}$=0.002; block 4, *V*=501, $p_{FDR}$=0.005) and outcome onset (block 1, *V*=499, $p_{FDR}$=0.008; block 2, *V*=485, $p_{FDR}$=0.011; block 3, *V*=447, $p_{FDR}$=0.037; block 4, *V*=575, $p_{FDR}=1.12\times10^{-4}$). Stronger block-level HPC RSA effects predicted more optimal behavioural (optimal paired-persist & rare-change) during character onset (Pearson's *r*=0.207, *P*=0.013), when participants had to decide whether to persist with or change the initial prediction. Correlation was marginally significant at the initial prediction phase (Pearson's *r*=0.151, *P*=0.071) and non-significant during the outcome phase (Pearson's *r*=0.077, *P*=0.360). Bar plots indicate the mean ± SEM. Each dot represents an individual. Fitted regression lines are plotted with 95% CI shading. **c-e.** Partial correlation RSA analyses (controlling for potential confounds) were performed separately for character onset (**c**), initial prediction onset (**d**), and outcome onset (**e**). See "RSA with partial correlation" in Methods for details. During the character onset, after regressing out the final goal prediction RDMs (see **g**), HPC still significantly represented the transition structure (*V*=532, *P*=0.009, after Holm–Bonferroni correction [HBC] for ROI number), with no comparable effects in other ROIs. During the initial prediction phase, after regressing out the initial goal prediction RDMs, the transition structure RSA effect of the anticipated characters remained significant in HPC (*V*=629, $P=1.17\times10^{-6}$, HBC), OFC (*V*=501, *P*=0.037, HBC), and lPFC (*V*=629, *P*=0.005, HBC). During the outcome phase, after regressing out the goal direction outcome RDMs, the RSA effects remained significant in HPC ($V$=611, $P$=1.64×$10^{-6}$, HBC) and vmPFC ($V$=520, $P$=0.016, HBC). **f.** Outcome-phase RSA (partial correlation) for correct trials only. HPC still showed significant encoding of transition structure ($V$=519, $P$=0.036), while vmPFC did not ($V$=428, $P$=0.139, HBC). The strength of this RSA effect positively correlated with participants' transition learning speed in the early learning period, as measured by regression of 'paired-persist' behaviour against pair history (Pearson's $r$=0.295, $P$=0.081). **g.** Partial correlation RSA of final goal predictions. Left panel: Example model RDM of final goal prediction, illustrating the dissimilarity between trials, coded as 0 for the same prediction and 1 for different predictions. RSA effects were computed after regressing out the dense transition structure RDM that does not omit self-comparison (see Fig. 4a and 'Representational similarity analysis (RSA)' in Methods). Among the tested ROIs, only OFC significantly encoded goal predictions during the character (final prediction) phase ($V$= 496, $P$=0.033, HBC). **h.** Transition structure RSA for correct trials. To control for trial count differences between correct and incorrect trials, we randomly subsampled correct trials to match the number of incorrect trials for each participant. This subsampling and RSA analysis were repeated 1,000 times to obtain stable group averages, the distribution of which was compared to a null distribution generated by shuffling model RDM labels. HPC showed significant RSA effects for correct trials ($P$<$10^{-16}$). ***$P$<0.001, **$P$<0.01, *$P$<0.05. n.s., not significant.

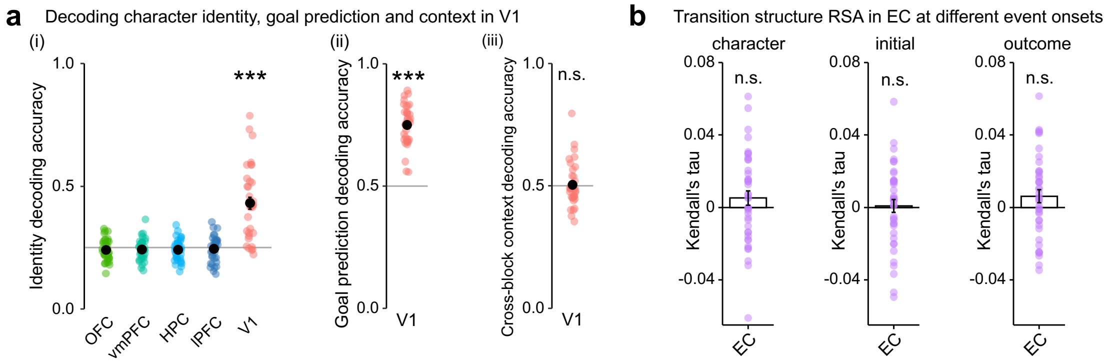


**Figure S6. Supplementary decoding and RSA analyses. a.** Decoding character identity, goal prediction, and context. (i) Only the primary visual cortex (V1) decoded the identities of four characters above chance ($t(35)=7.503$, $P=2.17\times10^{-8}$, HBC), while other ROIs showed were at chance level. (ii) V1 could also decode final goal predictions ($t(35)=18.175$, $P<10^{-16}$), given the visual features (left vs right arrows). (iii) But, unlike HPC, V1 was unable to decode the pairing structure (context decoding: $t(35)=0.256$, $P=0.777$), demonstrating specificity of HPC's representation for abstract structural knowledge beyond visual features. Each dot represents one individual, and black dots with error bars indicate mean ± SEM. **b.** No transition structure representation in EC. RSA results showed that the entorhinal cortex (EC), which is upstream of the HPC and was thought to provide contextual information, did not represent transition structure at all event onsets (character onset: $V=501$, $P=0.227$; initial prediction onset: $V=438$, $P=0.715$; outcome onset: $V=524$, $P=0.128$). ***$P<0.001$, n.s., not significant.

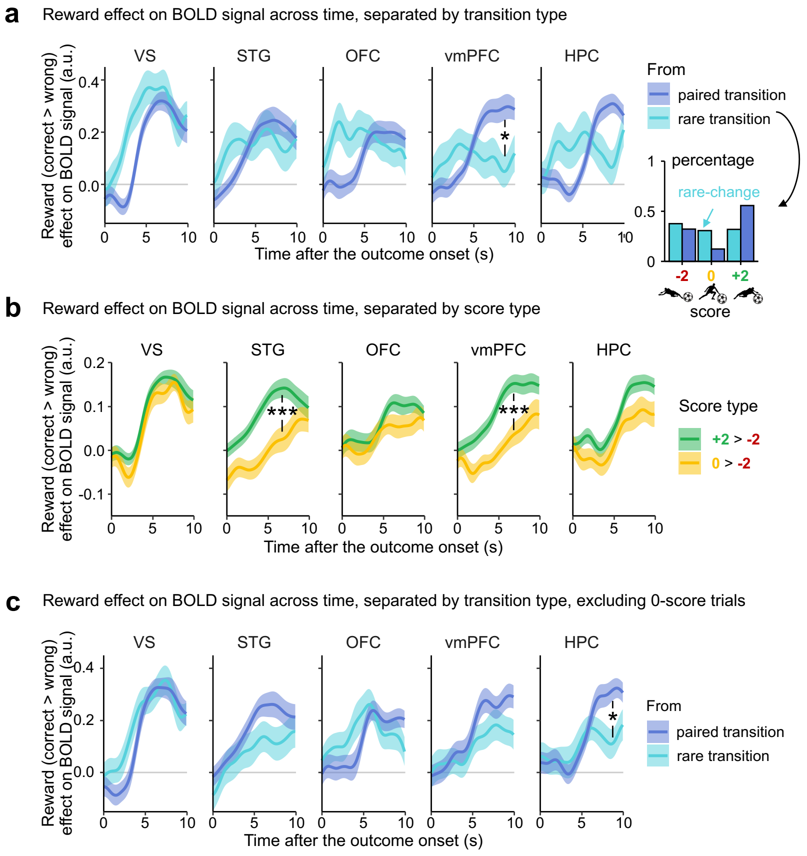


**Figure S7. Time-course analyses of reward effect on BOLD signal. a.** Time-course analyses of reward effects (correct > wrong) on BOLD signals in reward-related regions of interest (ROIs), plotted separately for paired and rare transitions. See "Time-course analysis" in Methods for details. Among the ROIs identified by whole-brain univariate analysis, only vmPFC showed significantly stronger reward-related activation during paired transitions compared to rare transitions (peak $V$=546, $p_{FDR}$=0.005, after HBC for ROI number: $p_{FDR}$=0.023). A similar effect in the HPC was significant before, but not after Bonferroni correction (peak $V$=516, $p_{FDR}$=0.030, after HBC: $p_{FDR}$=0.118). Notably, rare transitions prompted participants to change their initial predictions more frequently, resulting in a higher proportion of zero-score trials, whereas paired transitions had relatively more positive-score trials (inset). Thus, these differences may reflect value-related activation associated with the score of each trial. **b.** Time-course analyses separated by score types. To understand how different scores (values) influence reward-related activation, we separately examined BOLD responses for correct outcomes (positive, +2 points) versus neutral (0 points) outcomes, each compared to wrong outcomes (negative, -2 points). Significantly stronger activation for positive versus neutral outcomes was observed in STG (peak $V$=571, $p_{FDR}$=8.96×$10^{-4}$, HBC) and vmPFC (peak V=605, $p_{FDR}$=1.43×$10^{-4}$, HBC), highlighting their sensitivity to value-based reward signals. **c.** Time-course analyses of reward effects separated by transitions, after controlling for the influence of scores. After excluding zero-score, only HPC showed significantly stronger and sustained activation for correct outcomes following paired compared to rare transitions (peak $V$=538, $p_{FDR}$=0.0309, HBC), whereas the vmPFC effect was no longer significant. This result aligns with HPC's role in representing the transition structure, suggesting a reward-driven reactivation of learned transition knowledge. Ribbon plots show the group-averaged BOLD responses (beta estimates) across time (mean ± SEM). ***$P$<0.001, *$P$<0.05.

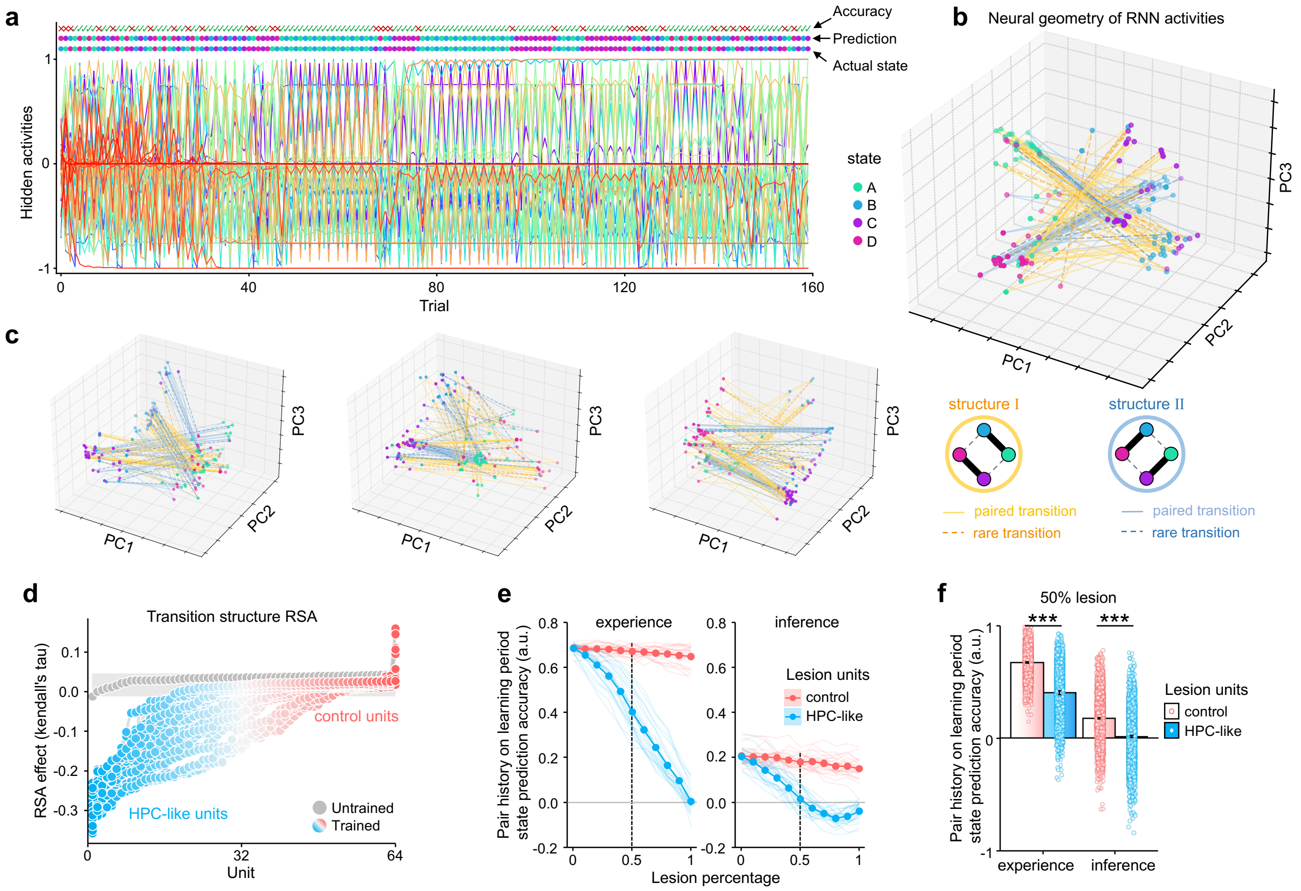


**Figure S8. Supplementary analyses of the neural network (RNN). a.** Example of unit activities within the RNN over one episode. RNN's state predictions, actual states, and prediction accuracies (match between prediction and actual state) across trials are shown on top. The RNN exhibited high prediction accuracy, except during rare transitions or reversals. Each coloured line in the lower panel indicates activity from a single unit across trials. Units encoded transitions differently from human HPC, maximising distinctions between paired characters, with larger activity changes within pairs. **b.** Example of neural geometry of RNN hidden activities (shown in **a**) based on principal component analysis (PCA; see "Principal component analysis" in Methods). PCA reduced the network's hidden-layer activity to a 3-dimensional neural space, with each dot representing neural activity on a single trial, coloured by the trial state. Solid lines represent paired transitions; dashed lines represent rare transitions. Two distinct pairing structures were used in the episode: structure I (AB & CD) transitions shown in orange/yellow, and structure II (AC & BD) transitions shown in light/dark blue. **c.** Additional examples of neural geometries from the same RNN across different episodes. **d.** Identification of transition-encoding (HPC-like) units via trial-level RSA in different RNNs trained with varied hyperparameters. Units encoding transition structure (blue, HPC-like) showed significantly different RSA values compared to baseline units from an untrained network (grey), or control units (red). Units from the same network are connected by grey lines. **e.** Effects of lesioning HPC-like units on transition learning across all trained networks. We separately lesioned HPC-like and control units in all the trained networks (See "Network lesion analysis" in Methods for details). As the percentage of randomly lesioned HPC-like units (blue) increases, all networks' use of both experience and inference (both $P<10^{-16}$) for predictions during the learning period declines. In contrast, lesioning control units (red) did not significantly impair learning. Thin transparent lines represent individual networks; thick solid lines indicate group means. **f**. A 50% lesioning of HPC-like units (indicated by dashed lines in **e**) significantly reduced the network's use of experience and inference, as shown in Fig. 6f. Bar plots indicate group mean ± SEM; Dots indicate individual simulation episodes from each network. ***$P<0.001$.

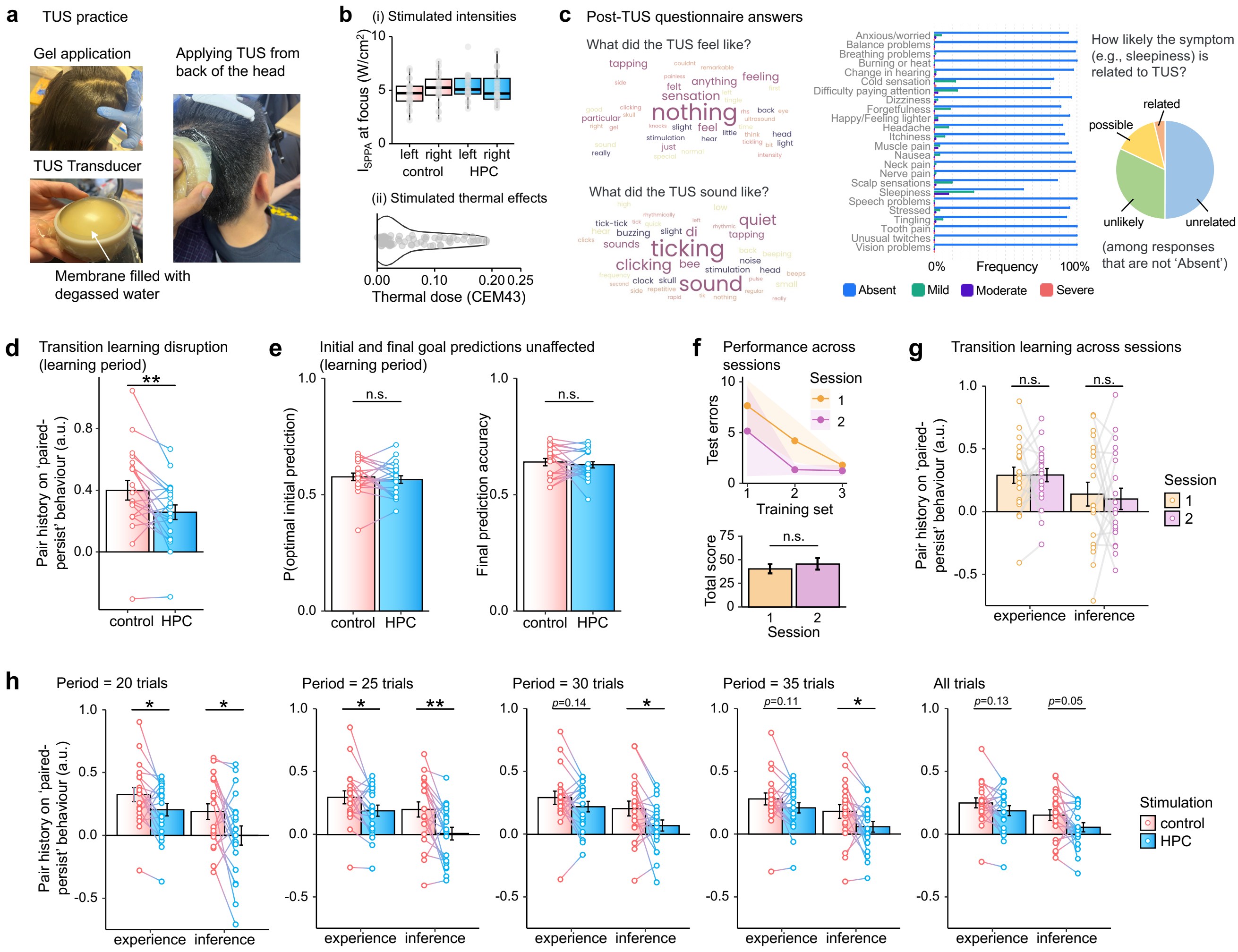


**Figure S9. Supplementary results from TUS experiment. a.** Photos illustrating the TUS procedure, including gel application and transducer placement (see "Applying TUS" in Methods). **b.** Intensity and thermal simulations. (i) Boxplots showing spatial peak pulse average intensity ($I_{SPPA}$) at the TUS focal point across participants, separately for the left and right hemispheres targeting HPC and control white matter regions. Intensities were approximately 5 W/cm² on average. (ii) The violin plot shows the distribution of thermal dose (CEM43) of all simulations (all smaller than 0.2). **c.** Post-TUS questionnaire results (see "Post-TUS questionnaire" in Methods). Text cloud visualises the most frequent answers to open questions related to sensations and sounds during TUS: most participants reported feeling "nothing" and hearing a "ticking" sound. Responses to potential adverse effects revealed that most symptoms were consistently rated as "Absent." For the few responses that reported symptoms (e.g., sleepiness), most participants judged these effects as unrelated or unlikely to be related to TUS, confirming the procedure's safety and tolerability. **d.** HPC-targeted TUS disrupted participants' transition learning compared to control stimulation during the learning period. Specifically, the effect of recent paired-transition history on optimal behaviour (paired-persist) was significantly reduced following HPC TUS (paired t-test: $t(19)=-2.913$, $P=0.009$). **e.** HPC stimulation did not significantly affect the optimality of initial predictions ($t(19)=0.720$, $P=0.480$) or the accuracy of final goal prediction ($t(19)=0.654$, $P=0.521$) during the learning period. **f.** Basic behavioural performance across the two TUS sessions. Participants learned the basic transition structure, reflected by a significant decline in their test errors across training sets. Two-way repeated ANOVA revealed significant training set effect ($F(2,38)=3.736$, $P=0.033$) with a marginal session effect: ($F(1,19)=3.513$, $P=0.076$), and no significant interaction between them ($F(2,38)=0.132$, $P=0.877$). There was no significant difference in the total scores participants earned between the two TUS sessions ($t(19) = 0.743$, $p = 0.467$). **g.** No systematic changes across sessions were found for the influence of experience ($t(19)=-0.021$, $P=0.983$) or inference ($t(19)=-0.331$, $P=0.744$), ruling out potential confounding session effects. **h.** HPC TUS disrupted the influence of experience and inference across different analysis windows (trial lengths). Significant effects were consistently observed for shorter periods (20 and 25 trials) after start or reversals of each block, while the effects weakened over longer periods, reflecting participants' ability to compensate for the disruption effect by accumulating knowledge over time. Inference-related disruptions remained more robust across all periods: 20-trial period: experience $t(19)=-2.441$, $P=0.025$, inference $t(19)=-2.527$, $P=0.02$; 25-trial period: experience $t(19)=-2.290$, $P=0.034$, inference $t(19)=-3.253$, $P=0.004$; 30-trial period: experience $t(19)=-1.537$, $P=0.141$, inference $t(19)=-2.490$, $P=0.022$; 35-trial period: experience $t(19)=-1.679$, $P=0.110$, inference $t(19)=-2.286$, $P=0.034$; 40-trial (all trials): experience $t(19)=-1.388$, $P=0.181$, inference $t(19)=-2.074$, $P=0.052$; Bar plots indicate group mean ± SEM. Dots connected by lines represent each participant's data. ***$P<0.001$, **$P<0.01$, *$P<0.05$. n.s., not significant.

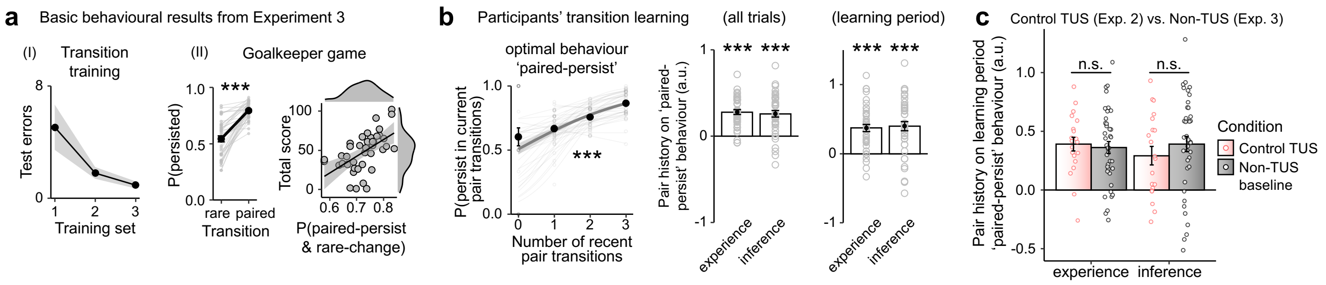


**Figure S10. Behavioural results from Experiment 3 (Behavioural-only).** Experiment 3 replicated the key behavioural findings from the fMRI experiment (Experiment 1), confirming the robustness of transition learning effects in an independent sample without neural measurements. **a.** Replication of basic behavioural performance. (i) During the Transition Training, participants implicitly learned the underlying transition structure, reflected by a gradual decline in their test errors across training sets ($F(2,78)=4.894$, $P=0.010$). Lines with shaded areas represent mean ± SEM. (ii) In the Goalkeeper Game, participants demonstrated optimal use of transition knowledge by persisting significantly more during paired transitions than during rare transitions (paired *t*-test: $t(39)=9.550$, $P=9.26\times10^{-12}$). The proportion of paired-persist & rare-change behaviour is positively correlated with their total score (Pearson's $r=0.553$, $P=1.31\times10^{-3}$). Fitted regression lines are plotted with 95% CI shading. Distributions of the data are shown on the edges of the plot. **b.** Regression analyses replicated key findings. Regression analyses (LM1, see Methods) confirmed that recent pair history significantly increased participants' likelihood of 'paired-persist' behaviour ($\beta\pm$CI=2.069±0.239, $P<10^{-16}$), and both the influences of same-pair experience ($\beta\pm$CI=2.015±0.219, $P<10^{-16}$) and other-pair inference ($\beta\pm$CI=1.766±0.228, $P=1.01\times10^{-14}$) were significant. When limiting the analysis to the learning period, the influence of both experience ($\beta\pm$CI=2.033±0.274, $P=1.18\times10^{-13}$) and inference ($\beta\pm$CI=2.120±0.325, $P=7.23\times10^{-11}$) remained robust. **c.** Consistency with the control TUS condition of the TUS experiment (Experiment 2). Participants' use of experience and inference in Exp. 3 (non-TUS baseline) closely matched those in the control TUS condition of Experiment 2 (two-sample t-test, experience: $t(45.134)=0.376$, $P=0.709$, Bayesian factor $BF=0.290$; inference: $t(44.04)=-0.982$, $P=0.332$, Bayesian factor $BF=0.394$). Bar plots indicate group mean ± SEM. Dots connected by lines represent each participant's data. ***$P<0.001$. n.s., not significant.
